# Unconstrained Plasticity Disrupts Memory Consolidation in a Mouse Model of Rett Syndrome

**DOI:** 10.64898/2026.01.29.702595

**Authors:** Tianrui Rui Zhang, Guanchun Li, Jeffrey C. Magee, Huda Y. Zoghbi, Sachin P. Vaidya

## Abstract

Memory impairment is a hallmark cognitive deficit in Rett syndrome (RTT). Yet, long-term memory deficits in RTT animal models remain poorly understood, largely due to the technical challenges inherent in tracking neural activity over extended periods. Here, we used longitudinal two-photon calcium imaging to follow the same population of hippocampal CA1 neurons as female RTT mice and their littermate controls formed cognitive maps of their environment during a spatial learning task. Neural representations in RTT mice were marked by excessive place cell (PC) activity, with individual PCs exhibiting pronounced instability across days. This disrupted single-cell stability propagated to the population level, resulting in unstable ensemble codes that poorly retained previously learned task information. Both excessive PC recruitment and instability could be attributed to a higher incidence of behavioral timescale synaptic plasticity (BTSP) in RTT mice. In wild-type littermates, place-cell consolidation across days is reflected by an increased likelihood of neuron-specific synaptic plasticity at the location of prior PC coding. This cellular mechanism of memory consolidation based on the location of BTSP was disrupted in RTT mice, where excessive and ectopic plasticity reduced PC stability, and degraded long-term stable representations. Backed by theoretical modeling, these results identify a plausible cellular and circuit-level mechanism underlying memory impairments in RTT mice and suggest principles that may be generalized to other neurological disorders involving memory deficits.

## Introduction

Rett syndrome (RTT), caused by loss-of-function mutations in the X-linked gene encoding the transcriptional regulator methyl-CpG binding protein 2 (MeCP2), is a leading genetic cause of intellectual disability in females^1,2^. Following an initial period of apparently normal development (6–18 months), affected individuals undergo developmental regression characterized by motor, linguistic, social, and cognitive impairments^2–4^. Female mice heterozygous for *Mecp2* (*Mecp2^+/-^*, hereafter referred to as RTT mice)^5^, which recapitulate core RTT phenotypes, exhibit markedly altered neuronal transcriptional profiles^6^ and structural deficits, such as reduced dendritic complexity and decreased dendritic spine density^7^, in the hippocampus, a brain region critical for learning and memory. Functionally, MeCP2 deficiency impairs Hebbian synaptic plasticity^8–11^, disrupts the excitation–inhibition balance and promotes hypersynchronous neural activity^7,12,13^. Although RTT mice exhibit learning and memory deficits in contextual fear conditioning and spatial memory tasks^7,11,13–15^, it remains unclear how these neurophysiological abnormalities affect the formation and consolidation of long-term memory.

To assess the emergence and stabilization of long-term memory in RTT mice, we tracked the evolution and persistence of place cells (PCs) in the hippocampal CA1 area during an experience-dependent task spanning multiple days. PCs are the hippocampal pyramidal neurons that encode specific locations in a given environment supporting navigation and memory-guided behavior^16–19^. With advances in longitudinal calcium imaging, PC dynamics now provide a powerful framework for dissecting mechanisms underlying long-term memory formation and consolidation in the hippocampus, an area long considered essential for episodic memory^20–30^. Growing evidence indicates that task-related learned information in the hippocampus is preserved in a subset of stable PCs that support subsequent memory retrieval^25,31–34^. At the cellular level, *in vivo* recordings of membrane potential and calcium dynamics implicate behavioral timescale synaptic plasticity (BTSP) as a key mechanism in PC formation in CA1 pyramidal neurons^34–41^. BTSP is a directed form of synaptic plasticity in which dendritic plateau potentials, driven by entorhinal inputs, potentiate recently active CA3–CA1 synapses, generating robust PC activity^42^. Recent work demonstrates that, when directed in a neuron- and location- specific manner, BTSP also serves as a primary mechanism for daily reconstitution of stable activity in PCs^34,41^. However, it remains unexplored how these synaptic plasticity mechanisms of long-term memory formation and consolidation are affected in neural circuits with MeCP2 deficiency.

## Results

### Elevated PC activity accompanies memory deficits in RTT mice

To characterize long-term memory and behavior-relevant neural dynamics in the mouse model of RTT, we performed longitudinal *in vivo* two-photon calcium imaging to record the activity of the same population of CA1 pyramidal neurons over seven consecutive days (see Methods), as head-fixed mice were engaged in a spatial learning task (Fig. 1a and Extended Data Fig. 1a,b). Female RTT mice and their wild-type (WT) littermates, maintained under controlled water access, were habituated to run on a 180-cm-long featureless treadmill belt. During habituation, a 10% sucrose solution was delivered pseudo-randomly at arbitrary locations along the blank belt, independent of the animals’ licking behavior. The final habituation session was designated as day 0 of the behavioral task. On day 1, mice were introduced to a novel 180-cm belt containing discrete sensory cues, with the reward location fixed at the 90-cm position throughout the subsequent 7-day training period. Reward delivery remained independent of licking behavior and did not influence the trial structure. (WT: *n* = 10 mice, 70 sessions, 100 ± 0.3 laps/session; RTT: *n* = 9 mice, 63 sessions, 99 ± 0.3 laps/session).

**Fig. 1.**
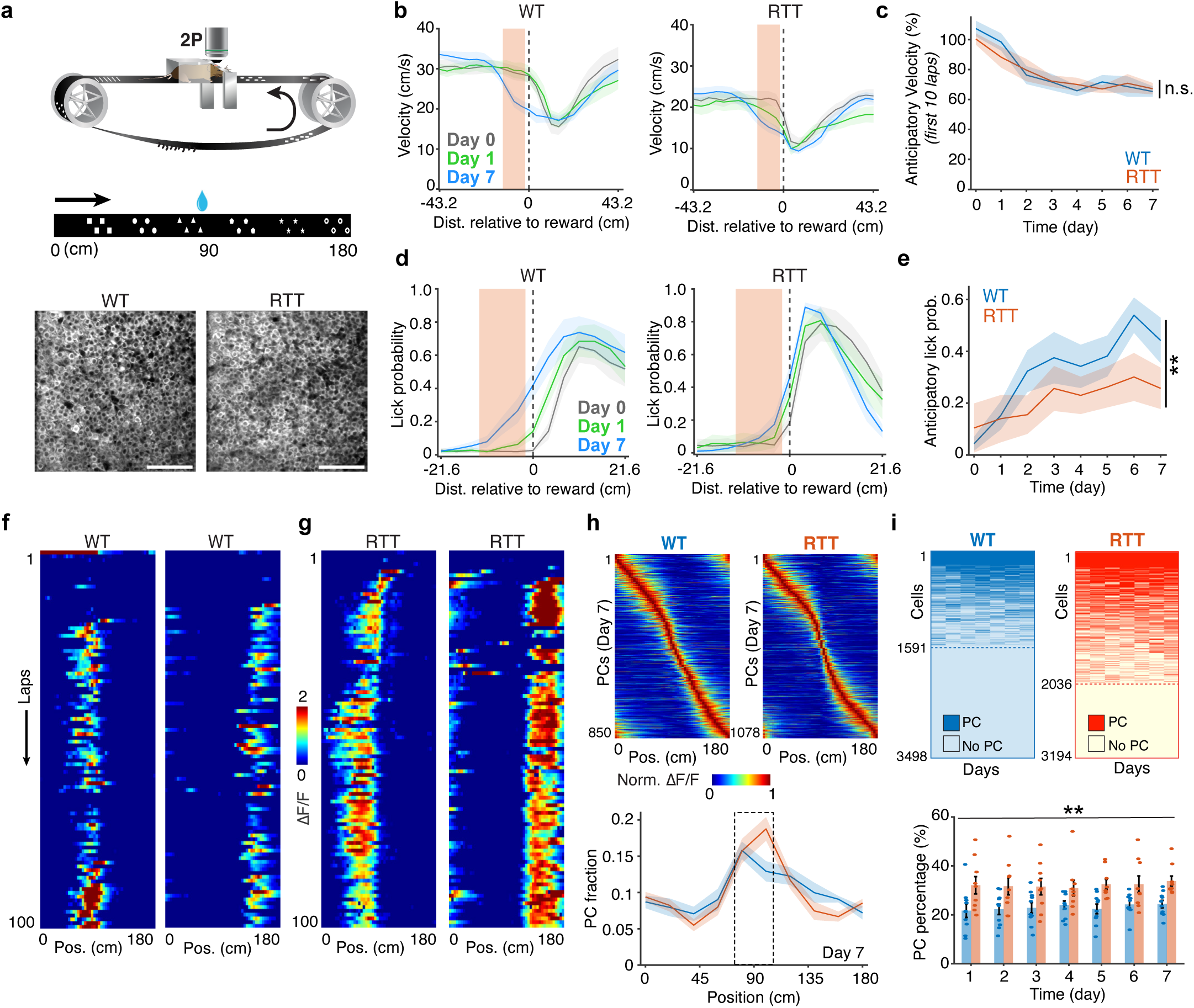
Elevated PC activity accompanies impaired task performance in RTT mice. **a,** Top: the experiment apparatus to record behavioral metrics and neuronal activity in head-fixed mice. Middle: the spatial learning task in which the mice learn water reward location on a 180-cm long linear treadmill enriched with sensory cues. Bottom: time-averaged images showing representative field of views (FOVs) of a WT mouse (left, RZ048) and an RTT mouse (right, RZ054) on day 1. The FOVs of these two animals on later days are shown in Extended Data Fig. 1. Scale bar, 100 µm. **b,** Average running velocity around the reward location in WT (left) and RTT (right) mice during the first 10 laps on day 0 (the last day of the training phase), day 1, and day 7. Orange boxes indicate the anticipatory zone, in which the velocity is measured for comparison in **c**. **c,** Change of running velocity in the anticipatory zone relative to that in a remote zone (see Methods) versus time. Linear mixed-effects model, time: *p* = 1.606 × 10^−8^; genotype × time interaction: *p* = 0.21495. n.s., genotype × time interaction *p* > 0.05. **d,** Lick probability around the reward location in WT (left) and RTT (right) mice during the entire sessions on day 0, day 1, and day 7. Orange boxes indicate the anticipatory zone, in which the lick probability is measured for comparison in **e**. **e,** Lick probability in the anticipatory zone versus time. Linear mixed-effects model, time: *p* = 0.003; genotype × time interaction: *p* = 0.007. **, genotype × time interaction *p* < 0.01. **f,g,** Representative PCs from WT (**f**) and RTT mice (**g**). Two different cells from separate recording sessions are presented for each genotype. **h,** Top: Peak-normalized activity (ΔF/F) of all PCs on day 7 in WT mice (left) and RTT mice (right). PCs are combined across animals and sorted by their PF locations. WT, *n* = 850 PCs; RTT, *n* = 1078 PCs. Bottom: Normalized distribution of PF locations on day 7. The box indicates the reward zone. Bin = 18 cm. Chi-squared test, WT: df = 9, *p* = 1.383 × 10^−9^; RTT: df = 9, *p* = 2.471 × 10^−32^. **i,** Top: PF activity across days in WT (left) and RTT (right) mice. Each row represents a CA1 pyramidal neuron. Each column represents a day. Dark colors indicate PF activity. Light colors indicate not exhibiting spatially modulated activity. Cells are combined across animals. WT, *n* = 3498 cells; RTT, *n* = 3194 cells. Bottom: Percentage of PCs among imaged CA1 pyramidal neurons across days. Two-way repeated-measures ANOVA, genotype: *p* = 0.009; time: *p* = 2.353 × 10^−13^. **, genotype *p* < 0.01. **b**–**e,h,i,** WT, *n* = 10 mice; RTT, *n* = 9 mice. Data are presented as mean ± SEM.

Both WT and RTT mice exhibited experience-dependent adaptive behavior, as evidenced by anticipatory slowing before the reward location during the first 10 laps of each session (Fig. 1b, c and Extended Data Fig. 2a). The progressive reduction in running velocity preceding the reward at session onset indicates that both groups acquired aspects of the task and retained some information about the reward location across sessions. In contrast, anticipatory licking before reward delivery—a behavioral measure of expectation and memory-guided reward anticipation—increased progressively in WT mice but remained significantly attenuated in RTT mice throughout training (Fig. 1d, e and Extended Data Fig. 2b,e,f). This deficit was not accompanied by alterations in post-reward licking metrics (Extended Data Fig. 2c–e,g), arguing against a generalized impairment in licking or motor execution. Previous studies in RTT mice have similarly demonstrated deficits in spatial learning and memory that are independent of motor coordination impairments^7,10,11^. Furthermore, anticipatory licking has been established as a robust readout of learned, memory-guided behavior in water-restricted mice, even when licking is not required to obtain reward^19,34,41,43,44^. Together, these findings indicate that, although RTT mice acquired aspects of the task reflected in adaptive running behavior, they exhibited a selective impairment in memory-guided reward anticipation compared with WT mice.

As the mice learned the task with repeated experience, we recorded the activity of 3,498 CA1 pyramidal neurons from 10 WT mice and 3,194 neurons from 9 RTT mice across seven days by imaging neurons that transgenetically expressed GCaMP6f (see Methods, Extended Data Fig. 3a).

We found significant differences in the activity level of the two cohorts. The calcium transient activity during locomotion was significantly elevated in RTT mice compared with WT littermates, as quantified by both event rate and normalized area under the curve (AUC/s) across all seven days (Extended Data Fig. 3b–e). These findings indicate that the CA1 region in RTT mice is generally hyperactive compared with littermate controls.

Despite diminished spatial memory, the CA1 pyramidal neurons in RTT mice formed robust PCs (Fig. 1f,g). Core properties of these PCs, including onset timing, reliability, place field (PF) width, amplitude, and spatial information content, were largely comparable to those in the WT littermates (Extended Data Fig. 4a–e). Importantly, in RTT mice, the PCs tiled the entire track with a preserved overrepresentation of the reward zone, indicating that CA1 circuits in RTT mice retain the capacity for behaviorally relevant and learning-efficient spatial coding (Fig. 1h, Extended Data Fig. 5a,b)^38,45–48^. Strikingly, RTT mice exhibited an approximately 1.4-fold higher proportion of active PCs per day compared with their littermate controls (Fig. 1i; 7-day animal-wise mean ± SEM: WT: 23.12% ± 1.65%; RTT: 32.14% ± 2.69%). Yet, the characteristic skewed distribution of PC propensity in the CA1 area, where most neurons remain inactive and a minority consistently form PCs across days, was mostly preserved in RTT mice (Extended Data Fig. 4f). Taken together, these observations suggest that, although the fundamental features of spatial coding remain intact in RTT mice, the cellular or circuit-level mechanisms governing PC formation may be disrupted, leading to substantially elevated PC activity in these animals.

### Increased incidence of BTSP in RTT mice is associated with elevated PC activity

To understand the mechanisms underlying elevated PC activity in RTT mice, we examined signatures of BTSP, the primary mechanism by which CA1 pyramidal neurons acquire spatial tuning^34–41^. Prior studies in the models of RTT have reported disruptions in NMDA receptor subunit expression^9^, dendritic spine density^7,49–53^ and inhibitory circuit function^7,12,13^, all of which could alter the probability of BTSP events.

In CA1 pyramidal neurons, BTSP is driven by dendritic plateau potentials occurring in the distal tuft, typically initiated by an interaction between direct entorhinal inputs and backpropagating action potentials arising from proximal CA3 inputs^42,54^. A single dendritic plateau can robustly potentiate CA3–CA1 synapses active within a seconds-long temporal window before and after the plateau. The asymmetric BTSP plasticity kernel in the CA1 region preferentially enhances inputs that occurred just before the plateau, giving rise to a characteristic predictive shift in the induced PFs in reference to the plateau location^35,37^.

To identify putative BTSP events, we applied the criteria combining a strong preceding calcium transient and a predictive shift in the subsequent spatial firing as a functional signature of BTSP (see Methods). This approach reliably captured Ca^2+^ events exhibiting hallmark features of BTSP (Fig. 2a,b): (1) abrupt onset of spatially tuned activity following a large, independently detected calcium event (Fig. 2c); (2) a predictive and spatially backward shift in the activity of the subsequent laps relative to the induction plateau (Fig. 2d); (3) a positive correlation between running velocity at the time of plateau induction and the width of the resulting PFs (Fig. 2e)^34,35,38,39,41^.

**Fig. 2.**
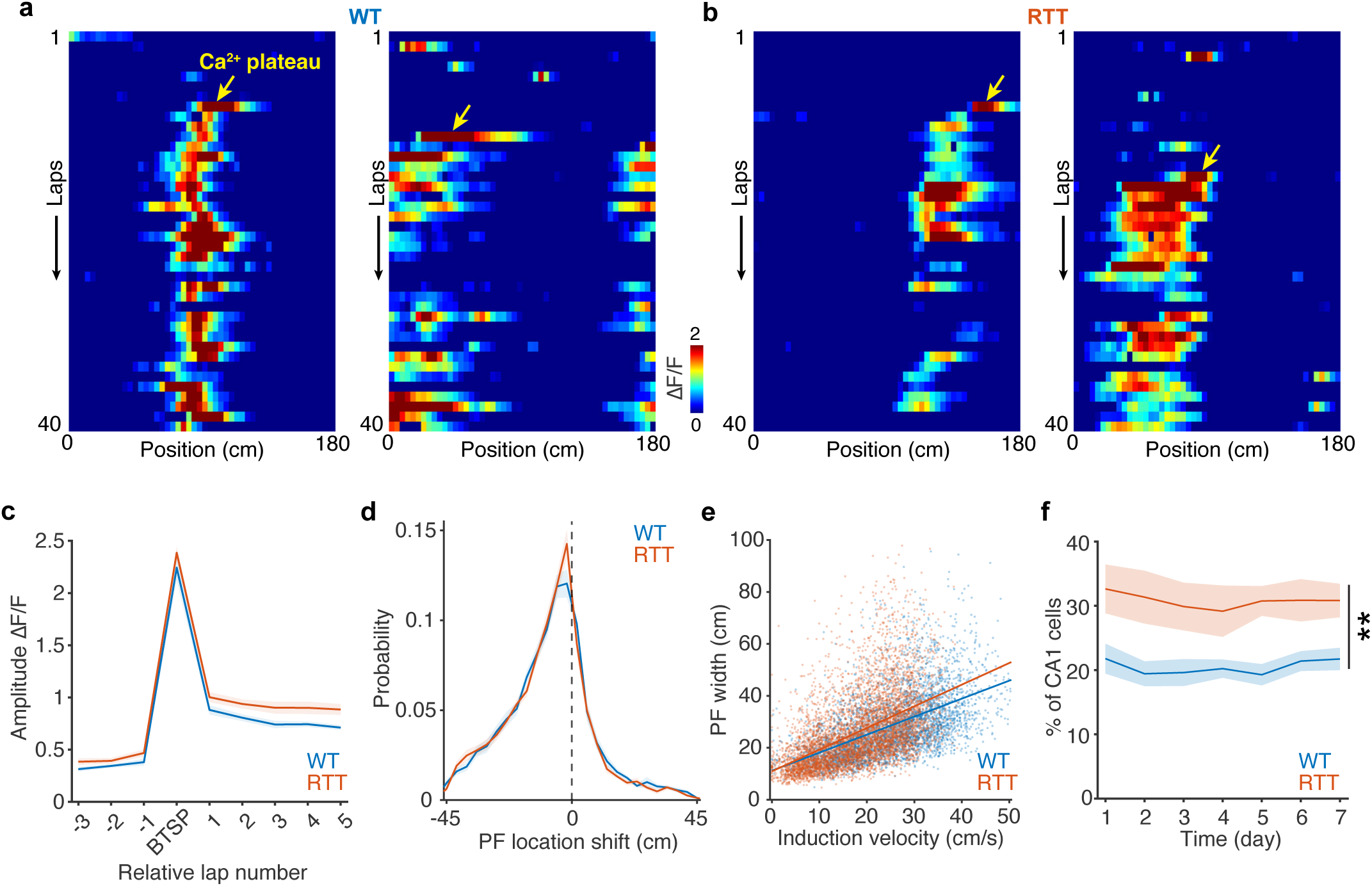
Increased incidence of BTSP in RTT mice. **a,b,** ΔF/F heatmaps across laps showing PF formation by BTSP in WT (**a**) and RTT (**b**) mice. Arrows indicate putative Ca^2+^ plateau potentials. Note the abrupt emergence of PF activity (measured in **c**) and the backward shift of subsequent PFs relative to the initial plateau location (measured in **d**). Two different cells from separate recording sessions are presented for each genotype. **c,** Average in-field ΔF/F (relative to the initial plateau location) across laps aligned to the BTSP induction lap. Data are from PCs on the first appearing day for a given PF location. BTSP events are combined across days. **d,** Normalized distribution of shifts between the initial plateau location and the PF location after the BTSP induction lap. BTSP events are combined across days. Bin = 3.6 cm. The dashed line indicates x = 0 cm. The median of the distributions in both genotypes is –5.4 cm. **e,** PF width as a function of the animal’s velocity during the initial BTSP induction. Each dot represents a PF with BTSP. BTSP events are combined across animals and days. WT: *n* = 4067 BTSP events; RTT, *n* = 5658 BTSP events. A linear regression model was fitted to the data. WT: R^2^ = 0.28825, *p* = 7.656 × 10^−302^, y = 0.69x + 11.18 (blue line). RTT: R^2^ = 0.25576, *p* < 0.001, y = 0.83x + 10.94 (orange line). **f,** Percentage of the cells having BTSP events among imaged CA1 pyramidal neurons across days. Two-way repeated-measures ANOVA with the Greenhouse-Geisser correction, genotype: *p* = 0.007; time: *p* = 7.145 × 10^−12^. **, genotype *p* < 0.01. **c,d,f,** WT, *n* = 10 mice; RTT, *n* = 9 mice. Data are presented as mean ± SEM.

Putative BTSP events were identified in over 70% of PCs in both groups (Extended Data Fig. 6a; 7-day animal-wise mean ± SEM: WT: 72.25% ± 2.17%; RTT: 77.04% ± 2.23%), with no detectable differences in the qualitative features of BTSP and plateaus between genotypes (Extended Data Fig. 6d–f). These features suggest that the core synaptic machinery underlying BTSP remains functionally intact in RTT mice. However, the overall incidence of BTSP significantly increased in RTT mice throughout the entire track, with the likelihood of BTSP induction approximately 1.5-fold greater than that in WT mice (Fig. 2f; 7-day animal-wise mean ± SEM: WT: 20.49% ± 1.59%; RTT: 30.77% ± 3.10%; Extended Data Fig. 6c). A strong correlation between the proportion of BTSP+ neurons and the proportion of PCs in both genotypes suggests that the increased incidence of BTSP events in RTT mice could account for a higher proportion of PCs observed above (Extended Data Fig. 6b). Taken together, these results imply that the regulatory mechanisms that constrain plateau potentials and synaptic plasticity may be impaired in RTT mice.

We further compared the coding properties between BTSP+ PCs and BTSP- PCs (Extended Data Fig. 7a–c). Within each genotype, BTSP- PCs exhibited later onset, lower reliability, reduced spatial information, lower activity amplitude, and narrower PF width than their BTSP+ counterparts (Extended Data Fig. 7d–h). Within each class of PCs, we observed no significant difference between WT and RTT mice (Extended Data Fig. 7d–h), and both classes of both genotypes showed overrepresentations of the reward zone (Extended Data Fig. 7i). We thus conclude that BTSP- PCs are systematically less precise and less robust. The properties of BTSP+ PCs reinforce the critical role of BTSP in establishing robust CA1 representations.

### Enhanced plasticity in RTT mice is accompanied by impaired stability of CA1 representations

A robust memory features not only appropriate initial encoding of environmental information, but also the persistence of these previously established neural representations. Central to this process is the delicate balance between plasticity and stability. While increased plasticity facilitates encoding new information, it can also render stored information susceptible to interference and overwriting, thereby compromising long-term memory retention^55–57^. By longitudinally recording the activity of the same CA1 neuronal population over extended periods (Fig. 3a,b), we were able to directly observe how this plasticity–stability trade-off was affected in the RTT hippocampus with higher incidence of plasticity.

**Fig. 3.**
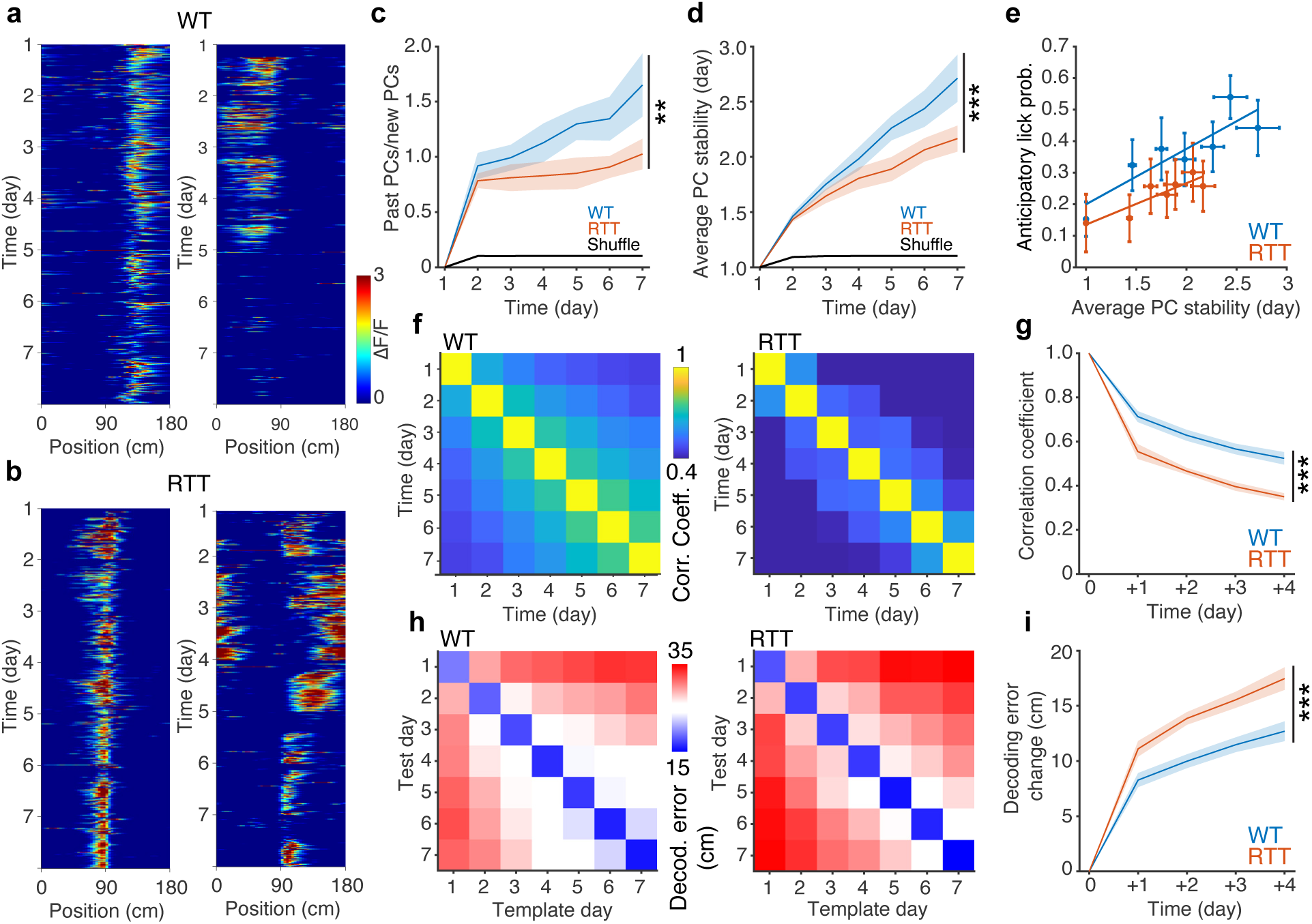
Long-term stability of CA1 spatial representations is impaired in RTT mice. **a,b,** ΔF/F heatmaps showing example cells from WT (**a**) and RTT (**b**) mice with varying PF activity through the 7-day experiment. ΔF/F of each lap is Gaussian filtered for visualization. **c,** Ratio of past PCs (maintaining previously established PFs) to new PCs (form PFs at new locations) versus time. The black line indicates data from a random process. Linear mixed-effects model, time: *p* = 4.563 × 10^−6^; genotype × time interaction: *p* = 0.005. **, genotype × time interaction *p* < 0.01. **d,** Average PC stability measuring the number of days during which the PC has maintained the same PF location versus time. The black line indicates data from a random process. Linear mixed-effects model, time: *p* = 8.634 × 10^−23^; genotype × time interaction: *p* = 1.370 × 10^−5^. ***, genotype × time interaction *p* < 0.001. **e,** Probability of anticipatory licks versus average PC stability. Each dot represents the probability of anticipatory licks and the average PC stability on an experiment day. A linear regression model was fitted to the data. WT: R^2^ = 0.76878, *p* = 0.0096, y = 0.1757x + 0.0239 (blue line). RTT: R^2^ = 0.78271, *p* = 0.008, y = 0.1315x + 0.0036 (orange line). **f,** Average population vector correlation matrices across animals in WT (left) and RTT (right) mice. **g,** Population vector correlation versus time increments (see Methods). Two-way repeated-measures ANOVA with the Greenhouse-Geisser correction (data from +1 day to +4 day), genotype: *p* = 5.871 × 10^−5^; time: *p* = 1.376 × 10^−16^. ***, genotype *p* < 0.001. **h,** Average decoding error matrices across animals in WT (left) and RTT (right) mice. **i,** Change of decoding errors relative to the same-day decoding versus time increments (see Methods). Two-way repeated-measures ANOVA with the Greenhouse-Geisser correction (data from +1 day to +4 day), genotype: *p* = 0.0006; time: *p* = 3.391 × 10^−15^. ***, genotype *p* < 0.001. **c**–**i,** WT, *n* = 10 mice; RTT, *n* = 9 mice. **c**–**e,g,i,** Data are presented as mean ± SEM.

To assess the stability of long-term memory representations in RTT mice, we analyzed both cellular- and population-level measures of PC stability across days. We first compared the ratio of the PCs that retained their spatial tuning from the previous days (past PCs, PF location shift ≤ ±30 cm) to those that formed PFs at a new location (new PCs, PF location shift > ±30 cm or the cells were silent on the prior day). This analysis revealed a pronounced and progressive divergence between genotypes over the 7-day learning period (Fig. 3c and Extended Data Fig. 8c). In WT mice, this ratio increased steadily with experience, pointing to the development of stable spatial representations. In contrast, RTT mice failed to show such accumulation, indicating a disruption in experience-dependent stabilization of PC activity.

At the level of individual neurons, we further computed the average stability of all PCs (number of days at the same location on the day of observation). On average, PCs in RTT mice were significantly less stable, suggesting they retained far less prior location information than those in WT controls, with the difference becoming more pronounced as the animals gained task experience (Fig. 3d). Importantly, the probability of accurate anticipatory licking was positively correlated with average PC stability across sessions (Fig. 3e), a relationship consistent with previous reports linking hippocampal representational stability to memory-guided behavior^19^.

Previous work suggests that the foundation of long-term memory in the hippocampus is supported by a minority population of highly stable, information-rich PCs that persist amidst a background of more transiently active cells^25,31,33,34^. The recruitment and accumulation of these highly stable cells, as a percentage of active PCs, was significantly reduced in RTT mice, further supporting the idea that hippocampal memory consolidation is impaired in the mouse model of RTT (Extended Data Fig. 8a,b,d).

Deficits in the long-term PC stability in RTT mice were further reflected in population-level measures of CA1 representations. Specifically, we found that cross-day population vector correlations were considerably diminished in RTT mice compared with WT controls, indicating less consistency in CA1 activity patterns over time (Fig. 3f,g). A linear decoder that was trained to predict the animal’s location on the track based on recorded neuronal activity (Extended Data Fig. 9a–d) confirmed the volatility of the spatial codes in RTT mice. The decoder error increased with longer intervals between the template and test day. The difference in this error, however, was significantly higher in RTT mice as compared with WT controls (Fig. 3h,i). This excessive decline in decoding accuracy further indicates a failure to retain stable neural representations with accumulating experience in RTT mice.

Together, these results suggest that despite enhanced BTSP and place-cell activity, RTT mice fail to develop a stable CA1 spatial representation over the same task experience. This instability is evident at both the cellular and population levels, pointing to an intrinsic deficit in place-cell stabilization in which excess plasticity undermines the persistence of spatial representations and, ultimately, long-term memory.

### Diminished spatial specificity of BTSP induction impairs PC stabilization in RTT mice

To elucidate the mechanisms underlying disrupted stabilization of PCs in RTT mice, we examined how individual PCs emerged and consolidated spatial information during repeated task exposures (Fig. 4a). The likelihood of a PC re-emerging at the same location on a given day increased sharply as a function of the PC’s prior history of encoding that location, providing direct evidence for an experience-dependent consolidation process at the single-cell level (Fig. 4b). Notably, the rate of this consolidation was modestly but significantly reduced in RTT mice compared with WT controls, indicating an impaired experience-dependent ability of maintaining stable PFs in RTT mice (Fig. 4b).

**Fig. 4.**
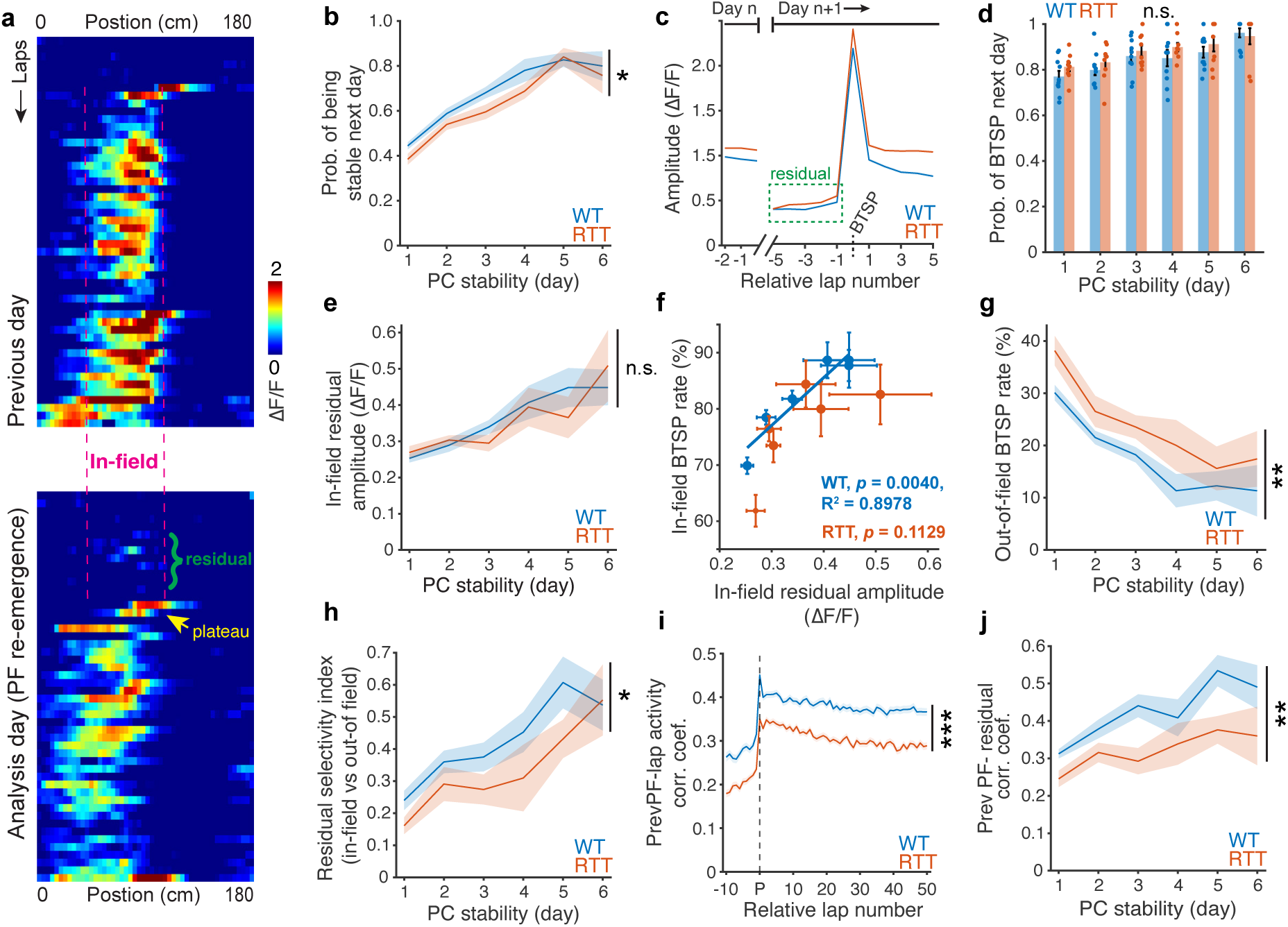
Diminished spatial specificity of BTSP induction impairs PC stabilization in RTT mice. **a,** A representative cell from WT mice showing the BTSP-induced PF reconstitution and the spatial specificity of pre-BTSP residual activity. Top: PC activity on the previous day; Bottom: PC activity on the following day. **b,** Probability of having a stable PF on the following day given the number of days for which a PF has been stable. Two-way repeated-measures ANOVA with the Greenhouse-Geisser correction, genotype: *p* = 0.038; time: *p* = 1.755 × 10^−20^. *, genotype *p* < 0.05. **c,** Lap-wise PF amplitude during re-emergence of spatially modulated activity. Data on day n show the last three firing laps (see Methods). Laps are aligned to the BTSP induction lap on day n+1 (1 ≤ n ≤ 6). Note the weak residual activity in the green box before a BTSP event reconstitutes active PF firing on the subsequent day. BTSP events are combined across animals and days. WT, *n* = 2614 BTSP events; RTT, *n* = 3368 BTSP events. **d,** Probability of having BTSP events on the following day given the number of days for which a PF has been stable. Two-way repeated-measures ANOVA with the Greenhouse-Geisser correction, genotype: *p* = 0.267; time: *p* = 2.768 × 10^−22^. n.s., genotype *p* > 0.05. **e,** Amplitude of in-field residual activity on the following day given the number of days for which a PF has been stable. Two-way repeated-measures ANOVA with the Greenhouse-Geisser correction, genotype: *p* = 0.837; time: *p* = 2.422 × 10^−11^. n.s., genotype *p* > 0.05. **f,** Probability of having in-field BTSP given the amplitude of in-field residual activity. Each dot represents the average in-field residual amplitude for a given number of days for which a PF has been stable and the probability of in-field BTSP event in that PC population. A linear regression model was fitted to the data. WT: R^2^ = 0.89782, *p* = 0.004, y = 85.0363x + 51.5592 (blue line). RTT: R^2^ = 0.50605, *p* = 0.113. There is statistical evidence that the two variables have linear relationship for WT mice but not for RTT mice, suggesting that increases in the amplitude of in-field residual do not effectively translate to enhanced in-field BTSP in RTT mice. **g,** Probability of having out-of-field BTSP on the following day given the number of days for which a PF has been stable. Two-way repeated-measures ANOVA with the Greenhouse-Geisser correction, genotype: *p* = 0.007; time: *p* = 5.243 × 10^−11^. **, genotype *p* < 0.01. **h,** Selectivity index of pre-BTSP residual activity (see Methods) on the following day given the number of days for which a PF has been stable. Two-way repeated-measures ANOVA with the Greenhouse-Geisser correction, genotype: *p* = 0.029; time: *p* = 7.957 × 10^−13^. *, genotype *p* < 0.05. **i,** Pearson’s correlation coefficient between lap-wise activity on following day with the PF activity on the previous day. BTSP events are combined across animals and days. “P” denotes the BTSP lap. WT, *n* = 2619 BTSP events; RTT, *n* = 3368 BTSP events. Two-sample *t*-test, p-values were adjusted for multiple comparisons using the Benjamini–Hochberg procedure (*q* = 0.05). ***, *p* < 0.001. **j,** Pearson’s correlation coefficient between average pre-BTSP activity on the following day with previous day’s PF activity, given the number of days for which a PF has been stable. Two-way repeated-measures ANOVA with the Greenhouse-Geisser correction, genotype: *p* = 0.006; time: *p* = 3.665 × 10^−13^. **, genotype *p* < 0.01. **b,d**–**h,j,** WT, *n* = 10 mice; RTT, *n* = 9 mice. **b**–**j,** Data are presented as mean ± SEM.

To understand this impaired memory consolidation in RTT mice, we analyzed the dynamics of PF re-emergence, in which PFs reappeared at the previous location (*in-field*) or shifted to new positions (*out-of-field*) on the subsequent day. Consistent with previous reports, prior PF activity substantially decayed overnight, leaving only faint traces, if any, at the beginning of the next session (Fig. 4c). PF re-emergence was typically accompanied by a *de novo* BTSP event that reinstated spatially modulated activity. This phenomenon occurred in both cohorts where over 80% of re-emerging PFs were reconstituted by a plateau-driven BTSP event (Fig. 4c,d; animal-wise mean ± SEM: WT: 85.26% ± 1.29%, RTT: 88.05% ± 1.21%, all subsequent sessions combined, WT, n = 60 sessions, RTT, n = 54 sessions).

To determine why some reconstituting plateau potentials occurred within the original place-field location, producing a stable PF, whereas others occurred outside it, yielding an unstable PF, we examined the weak residual activity present at the former place-field location prior to the initial BTSP event. We refer to this activity as the *in-field residual*. The amplitude of the in-field residual increased with the spatial stability of the preceding PF in both groups (Fig. 4e). This relationship suggests a cellular consolidation mechanism that biases the reconstitution of the PF toward its previous location by providing location-specific depolarization which enhances the likelihood of BTSP at the given location. This mechanism remained largely unaffected by MeCP2 deficiency in RTT mice.

Consistent with this idea, in WT mice, the probability of in-field BTSP events strongly correlated with the experience-dependent increase in the in-field residual amplitude. Unexpectedly, however, RTT mice did not exhibit such a similar increase in the in-field BTSP probability despite showing a comparable rise in in-field residual amplitude (Fig. 4f). This impairment was further reflected by a higher probability of out-of-field BTSP events in RTT mice compared with WT controls, especially when the stability of the prior PF was considered (Fig. 4g). Together, these findings indicate that in RTT mice, higher in-field activity does not translate into an increased likelihood of in-field BTSP as effectively as in WT mice.

We next examined why the experience-dependent increase in in-field residual activity failed to confer strong spatial specificity of BTSP induction in RTT mice. To this end, we quantified a simple selectivity index that compared the in-field residual activity with its corresponding out-of-field pre-BTSP activity. This residual selectivity index revealed that, despite a similar increase in in-field residual amplitude, the contrast between in-field and out-of-field activity was reduced in RTT mice, particularly during early stages of consolidation (Fig. 4h). This result aligns with our earlier observation, that activity levels were markedly higher in RTT mice than in WT mice, perhaps obscuring the weak spatially tuned activity (Extended Data Fig. 3b–e). Lap-wise activity correlations with the previous PF activity further supported these observations, where the pre-BTSP activity in RTT mice showed a lower correlation compared with WT mice (Fig. 4i,j). Notably, the reduced correlation with the previous PF persisted even after the BTSP event, indicating greater PC displacement and instability as observed above (Fig. 4i).

Taken together, these findings identify a cellular substrate for memory consolidation in terms of an experience-dependent increase in weak residual activity at the location of the prior PF. The in-field residual activity would bias the subsequent BTSP event toward the previously active PF location, thereby increasing the likelihood of re-forming a stable PF. Multiple lines of evidence indicate that this experience-dependent consolidation mechanism is defective in RTT mice, producing an inaccurate location of BTSP induction and, consequently, impairing PC stability and memory retrieval.

### A computational model with distorted BTSP localization recapitulates PC dynamics in RTT mice

To understand how misdirected BTSP driven by noisy residual activity impacts long-term memory consolidation, we modeled the process of PC stabilization over the course of one week (see Methods, Fig. 5a,b and Extended Data Fig. 10a–d). In the model, the probability of BTSP and consequently PF formation at a given location was primarily determined by weak residual depolarization preceding the BTSP event. This pre-BTSP depolarization at a given location was reflective of previously active, resilient synaptic weights, such that locations with a history of sustained PF activity exhibited reduced overnight decay and elevated residual depolarization (Fig. 5a,b and Fig. 4c). This modeling framework recapitulated the experimentally observed increase in *in-field residual* activity with spatial stability (Fig. 5c and 4e). The weak pre-BTSP depolarization was also influenced by spontaneous network activity that was present across all locations and neurons.

**Fig. 5.**
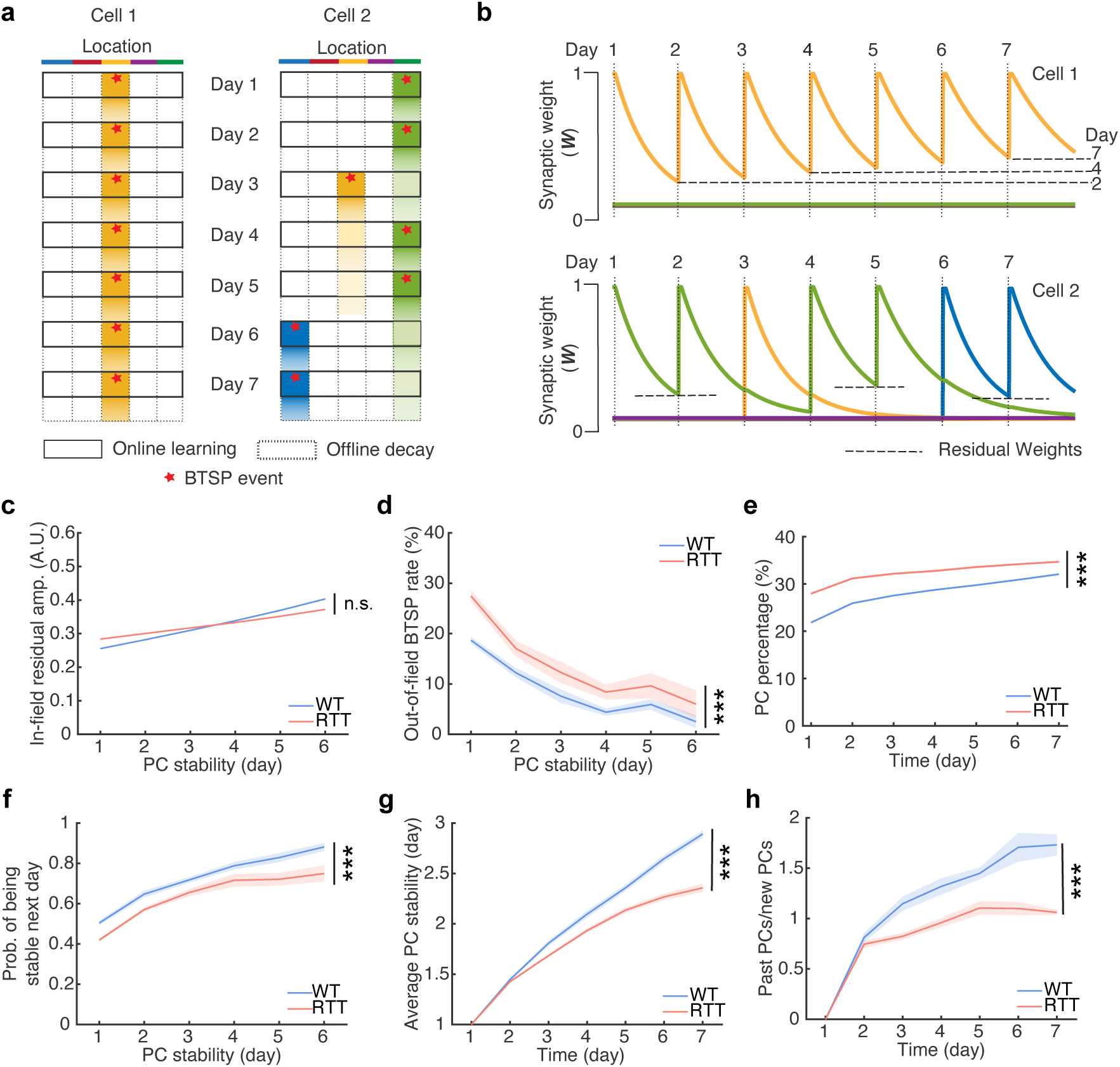
A computational model with altered residual selectivity recapitulates RTT PC dynamics. **a,b,** Two cells exemplifying the PC dynamics model. Cell 1, WT-like condition, maintain one PF location stably; Cell 2, RTT-like condition, change PF locations. The synapses encoding a given location are potentiated daily by BTSP during the online learning phase and their weights decay during the offline period. The colors indicate five distinct PF locations in an environment. The darker color depicts higher synaptic weights in **a**. Note the history-dependent enhancement of residual synaptic weights in **b**. **c**, The in-field residual amplitude stays comparable between the WT- and RTT-like conditions in this model. Two-way repeated-measures ANOVA with the Greenhouse-Geisser correction, genotype: *p* = 0.9. n.s., genotype *p* > 0.05. **d,e,f,** The PF dynamics model recapitulates the out-of-field BTSP probability (**d**), percentage of PCs (**e**), and experience-dependent PC stabilization (**f**) under the WT- and RTT-like conditions. Two-way repeated-measures ANOVA with the Greenhouse-Geisser correction. **d**, genotype: *p* = 1 × 10^−7^. **e**, genotype: *p* = 7 × 10^−12^. **f**, genotype: *p* = 6 × 10^−8^. ***, genotype *p* < 0.001. **g,h,** The PF dynamics model recapitulates the development of stable CA1 representations with task experience and the divergent responses between the WT- and RTT-like conditions. **g**, Ratio of past PCs (maintaining previously established PFs) to new PCs (form PFs at new locations). Two-way repeated-measures ANOVA with the Greenhouse-Geisser correction, genotype: *p* = 1 × 10^−6^. ***, genotype *p* < 0.001. **h**, Average PC stability measuring the number of days during which the PC has maintained the same PF location. Two-way repeated-measures ANOVA with the Greenhouse-Geisser correction, genotype: *p* = 4 × 10^−8^. ***, genotype *p* < 0.001. **c**–**h,** WT-like condition, *n* = 10 simulations; RTT-like condition, *n* = 10 simulations. Data are presented as mean ± SEM. The error bars in **c** and **e** are not visible because of the low variance.

To simulate RTT-like conditions, we increased the non-specific spontaneous activity (see Methods), a manipulation that may reflect impaired inhibitory control or heightened excitability in upstream circuits in RTT animals^13,59^, and is consistent with our experimental observations (Extended Data Fig. 3b–e). This single manipulation markedly reduced the residual selectivity index under RTT-like conditions, recapitulating the experimentally observed increase in noisy residual activity with prior PF activity and providing a mechanistic framework to distort the experience-dependent spatial specificity of BTSP (Extended Data Fig. 10e). The reduction in residual selectivity resulted in a significant proportional increase in the displacement of BTSP events away from previous PF locations, quantified as an elevated probability of *out-of-field* events in cells exhibiting PFs on two consecutive days, irrespective of the stability of the preceding PFs (Fig. 5d and 4g). In addition, the introduction of spontaneous activity increased the overall incidence of plasticity, accounting for the higher fraction of PCs observed under RTT-like conditions and recapitulating another key experimental finding (Fig. 5e and 1i).

A central argument in our findings is that excessive and misdirected BTSP, likely arising from impaired regulatory mechanisms of synaptic plasticity, can disrupt memory consolidation at the level of single cells and, in turn, compromise experience-dependent learning and the stability of population representations. When we assessed consolidation rates in single cells within the model incorporating distorted BTSP targeting and elevated plasticity, we observed a significant reduction in single-cell stabilization rates across all stages of consolidation, closely mirroring the deficits measured in RTT animals (Fig. 5f and 4b). Although modest at the single-cell level, these differences accumulated over time to produce pronounced impairments in population-level stability. Specifically, the progressive reduction in the daily accrual of stable PCs was reflected in both a significant decrease in average single-cell stability and reduced fraction of past PCs, thereby recapitulating the diminished stability of CA1 representations observed experimentally in RTT mice (Fig. 5g,h and 3c,d).

Given the mosaic cellular environment in female *Mecp2^+/−^*mice, we also simulated RTT-like conditions by increasing non-specific spontaneous activity in only 50% of neurons that mimicked MeCP2-lacking cells. This manipulation in a subpopulation of cells recapitulated all the major phenotype from experiments and the previous simulation that introduced non-specific spontaneous activity in all the cells (Extended Data Fig. 11a–f). While this simple model does not distinguish the biophysical source of the increased spontaneous activity — impaired inhibitory control within CA1 or hyperexcitability in upstream regions — a single manipulation that reduces the prominence of residual activity was sufficient to disrupt place cell stabilization in the simulations. This suggests that, in principle, elevated non-specific activity is sufficient to compromise the stabilization of CA1 representations and, ultimately, long-term memory consolidation.

## Discussion

In this study, we characterized PC formation and stability as a model for long-term memory in RTT mice learning a spatial navigation task over multiple days. Despite well-documented synaptic, cellular, and circuit-level impairments in RTT mice, we find that RTT mice form robust PCs similar to WT littermates. However, RTT mice exhibit a ∼40% increase in the number of PCs, indicating disrupted mechanisms controlling PC induction. Notably, these PCs are predominantly generated through BTSP, a plasticity mechanism that remains intact but insufficiently regulated, despite impaired Hebbian plasticity in RTT mice. Over extended period, RTT mice show deficits in spatial memory-guided behavior, coinciding with an abnormally high turnover of highly labile PCs, suggesting that excessive instability, rather than failure to initially form PFs, may underlie long-term memory impairments in RTT. In WT mice, stable PFs appear to arise from a directed BTSP event targeted to the location of prior PC activity, likely guided by weak pre-existing firing at that site. Our experimental findings, supported by the modeling of PC dynamics, indicate that this process of experience-dependent directed plasticity is disrupted in RTT mice, resulting in impaired stabilization of PCs, and ultimately, deficits in spatial memory.

### Dysregulated BTSP arises from weakened dendritic inhibition in the CA1 microcircuit

Female RTT mice used in this study (*Mecp2^+/-^*;*Thy1*-GCaMP6f) have a random mosaic expression for the loss-of-function *Mecp2* allele, causing MeCP2 deficiency in ∼50% of neurons and, replicating the phenotype of human RTT^49^. The hippocampal circuit and its impairment of memory up to 24 hours have been well characterized in this mosaic model, particularly in the context of contextual fear conditioning and spatial memory tasks^7,11,13–15,49^. Kee et al. recorded hippocampal CA1 activity using tetrodes in freely moving RTT mice and reported that PC refinement with experience, in terms of width and spatial information, was impaired in these mice^15^. However, in that study, mechanisms of PC formation and extended stability remain unexplored due to the technical limitation of extracellular electrophysiology. Using two-photon microscopy to access large neuronal populations, we find that disrupted regulation of BTSP is responsible for excessive PC activity in this model. BTSP is a novel form of synaptic plasticity directed by excitatory inputs from the entorhinal cortex layer 3 (EC3), which provides a *target signal* for hippocampal representations of a given environment. EC3 activity, which is conveyed to the distal tuft of CA1 pyramidal neurons, increases the likelihood of dendritic plateau potentials that enable BTSP^38,42,60^. It is proposed that this plasticity is constrained by a feedback inhibitory loop, mediated by *oriens lacunosum-moleculare* (OLM) interneurons^61^. These interneurons integrate the overall activity of CA1 pyramidal neurons and provide corresponding dendritic inhibition that may limit dendritic plateau potential initiation when there is sufficient PC activity. Optogenetically silencing OLM interneurons during spatial learning resulted in an increased PC density in the opto-zone^61^, reminiscent of the enlarged population of PCs in RTT mice. In those animals, this OLM feedback circuit is weakened, largely due to poor recruitment of MeCP2-deficient OLM cells^13^. Because this feedback signal does not discriminate MeCP2-positive or MeCP2-negative CA1 pyramidal cells, its reduction likely affects the entire population of CA1 excitatory neurons. Previous work has linked this impaired feedback circuit to hypersynchronous CA1 activity during quiescent, non-locomotor states, with potential consequences for the refinement of contextual representations, particularly in fear conditioning paradigms^12,13,15,61^. Here, we propose a novel implication of the diminished dendritic inhibition that may alter the homeostatic balance by upregulating plasticity during awake behavior to distort PC coding in the hippocampus by increasing PC activity which is unstable in nature.

### Ectopic activity misdirects the plasticity underlying memory consolidation

One consequence of a highly plastic system is disrupted memory stability. In what is known as the stability–plasticity conundrum, theoretical models predict that highly plastic networks impair memory permanence owing to overwriting of previously stored memories during continual learning^55–57^. A proposed solution to this problem is the selective stabilization of reinforced memory units that are resistant to modification and thus protected from interference^57,62,63^. Recent work has identified signatures of such a *cascade* of stability within the hippocampus, where a subset of PCs, encoding salient features of the environment, become increasingly stable, forming an expanding core of task-relevant, learned representations^34^. In this study, we find that the process of single cell stabilization is impaired in RTT mice with excessive plasticity leading to poor preservation of learned information over time. Unconstrained plasticity in RTT hippocampus, when contrasted with littermate controls, thus provides a direct demonstration of the theorized disturbed stability–plasticity balance in an active biological circuit.

A closer look into the process of single PC stabilization in the hippocampus suggests that stability is not solely determined by the permanent modification of active synaptic weights. Instead, it is more intricately achieved by an increased probability of directed plasticity that potentiates weakened or new synapses to reconstitute PC activity at a specific location^34,41,58^. The mechanisms underlying this directed plasticity at the same location day after day, which are at the heart of memory consolidation, remain an active area of research. In this study, we present evidence that the subtle but notable residual activity, at the location of the previous PF, may be instrumental in directing the BTSP event which induces a stable PF at the correct location. As a signature of memory consolidation, this signal becomes stronger with experience, perhaps reflective of the aforementioned stability cascade where an increasing subset of strong synapses are more resistant to overnight decay. In RTT mice, this cellular mechanism of memory consolidation appears to be hijacked by ectopic activity that misdirects BTSP, compromising the stability of CA1 representations. This ectopic activity may be a result of higher spontaneous activity, perhaps an outcome of reduced inhibition in hippocampal CA1 microcircuits. It is also possible that it may arise from the upstream CA3 region, which in RTT mice is known to be hyperexcitable with increased spontaneous firing^59^.

Overall, the evidence provided here supports an alternative theory of long-term memory impairments in RTT mice, centered around dysregulated plasticity mechanisms. While prevailing theories of memory impairments in these mice focus on hypersynchronous activity and its potential interference with classic models of *offline* consolidation, our novel theory highlights disruption of active *online* cellular mechanisms of memory consolidation during awake locomotion. Ultimately, by comparing these findings from the mouse model of RTT and their littermate controls, we hope to gain insights into memory consolidation in healthy neural circuits as well as the malfunctions that characterize neuronal pathologies.

## Methods

All experimental procedures were approved by the Institutional Animal Care and Use Committee (IACUC) of Baylor College of Medicine (protocol AN-1013 and AN-7734).

### Animals

The data were collected from 10 female *Mecp2^+/+^*;*Thy1*-GCaMP6f (WT) mice (14–18 weeks) and nine female *Mecp2^+/-^*;*Thy1*-GCaMP6f (RTT) mice (14–16 weeks). They were F1 offspring generated by crossing female heterozygous *Mecp2^+/-^* mice^5^ (the Jackson Laboratory, 003890) maintained on C57BL/6J background with male mice homozygous for *Thy1*-GCaMP6f^64^ (line GP5.17, the Jackson Laboratory, 025393). Because female *Mecp2^+/-^* mice are often poor mothers, their litters were fostered by lactating female FVB/NJ mice (the Jackson Laboratory, 001800) that had given birth within 5 days of the birth of pups from *Mecp2^+/-^* mice until weaning. Animals were group-housed on a 14-h light/10-h dark cycle (lights on at 6 am and lights off at 8 pm) at 21 ℃ and 30–70% humidity, with standard chow and water provided *ad libitum*. The mice used for experiments were transferred to the Magee laboratory satellite facility at least two weeks before surgery (see below), where they were housed under a reversed 12-h dark/12-h light cycle (lights off at 9 am and lights on at 9 pm).

### Surgery

All surgical procedures were performed in mice between 10 weeks and 13 weeks under deep anesthesia with ∼1.8% isoflurane as previously described^34,38,41,65^. After locally applying topical anesthetic, the scalp was removed, and the skull was cleaned and leveled. A 3-mm-diameter craniotomy centered at 2.1 mm posterior from bregma and 2.0 mm from the midline (right hemisphere) was made above the right hippocampal CA1 area. Then, the dura was removed with forceps, and the overlying cortical tissue within the craniotomy was slowly aspirated using a blunt needle (CML Supply, 901-23-100 and 901-26-050) under constant irrigation with sterile normal saline. Once the external capsule was exposed, a cannula (3 mm in diameter and 1.95 mm in height) with a coverslip (CS-3R, Warner Instruments) glued onto the bottom using UV-curable optical adhesive (Norland Products, NOA81) was inserted and cemented to the skull. Finally, we attached a custom-made titanium head bar to the skull parallel to the plane of the imaging window using dental acrylic (Ortho-Jet, Lang Dental).

### Behavioral training and task

A 180-cm velvet fabric belt was used to train the animals to run and perform subsequent experiments. Mice were head-fixed to a custom-fabricated stainless-steel head bar holder. The belt was self-propelled by water-restricted mice, and the animal’s position and velocity were measured by a rotary encoder coupled to an Arduino-based microcontroller. All behavioral variables (position, velocity, lap markers, reward delivery, and licks) were digitized at 10 kHz via a PCIe-6343, X series DAQ system (National Instruments) using WaveSurfer software (wavesurfer.janelia.org). The position signal was interfaced with a behavior control system using a Bpod module (Sanworks) that controlled reward delivery (10% sucrose solution) through a solenoid valve (quiet operation, the Lee Company).

Mice were given a recovery period of at least seven days before further behavioral training. Following that, mice were water-restricted (1.5 mL per day) to motivate running on the treadmill. The behavioral training started with habituation to the experimenter, treadmill, and water rewards for about 15 min per day for at least nine days. The mice were then head-fixed and trained to run for water rewards delivered at random locations on a blank belt with no sensory cues. Meanwhile, they were accustomed to two-photon imaging during the later part of the training regimen. The mice would be introduced to the experimental task if they could run 100 laps (180 cm per lap) in 1 h on two successive days while the two-photon microscope was imaging the hippocampus. The animals that could not meet this criterion within 10 days would not be used for further experiments. The last day of the training phase was designated as day 0 (Fig. 1b–e).

On day 1 of the experimental task, the mice were introduced to a new belt containing six distinct sensory cues, each occupying 15 cm. The reward delivery location was fixed at 90 cm. The mice performed one session per day, and they were introduced to the same belt and task for seven consecutive days. On day 1, the WT mice were 14–18 weeks old, and the RTT mice were 14–16 weeks old.

The reward delivery was controlled by a quiet operation solenoid valve (LHQ series, The Lee Company) located away from the lick port to eliminate any potential auditory cues associated with reward delivery. Licks were detected by fiber optic sensors (FX-300 series, Panasonic) with custom fabrication. Lick detection did not affect the trial structure, since rewards are independent of licking.

### Longitudinal two-photon imaging

All *in vivo* Ca^2+^ imaging was performed in the dark using a custom-built two-photon microscope (Janelia MIMMS design). Transgenically expressed GCaMP6f was excited at 920 nm (typically 20–60 mW) by a Ti:Sapphire laser (Chameleon Ultra II, Coherent) and imaged through a 16X, 0.8-NA objective (Nikon). Emission light passed through a 565 DCXR dichroic filter (Chroma) and was detected by GaAsP photomultiplier tubes (11706P-40SEL, Hamamatsu). Images (512 × 512 pixels) were acquired at 30 Hz using ScanImage software (Vidrio Technologies). For each mouse, a reference field of view (FOV) (∼353 × 353 µm) was chosen before day 1 and then registered and imaged daily. Experiments would be terminated if FOVs on subsequent days showed substantial changes.

### Data analysis

#### Quantification of behavioral data

All analyses of behavioral metrics were conducted using custom code in MATLAB (version 2024a). Spatial profiles of running velocity, lick probability, number of licks, and dwell time were generated by dividing the 180-cm track into 50 bins (3.6 cm per bin) and aligning all behavioral measures to the reward location, defined as bin 25. For each spatial bin, the average velocity was calculated as the mean of all the recorded velocity that exceeded 2 cm/s. The velocity change r_i_ in each anticipatory zone (bin 22–24) was calculated as:

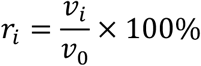

v_0_ is the average velocity of bin 1–6 the designated remote area. v_i_ is the average velocity of bin *i* (*i =* 22,23,24). The velocity change in the anticipatory zone was the mean of r_i_ of bin 22–24. Only the first 10 laps of each session were included to plot velocity traces and quantify the velocity change.

When plotting the lick probability traces, for a given spatial bin, if there was at least one lick in the bin during a lap, this lap would be considered as a ‘lick trial’ (Fig. 1d). When calculating the anticipatory lick probability, if there was at least one lick within 10.8 cm before the reward location during a lap, this lap would be considered as a ‘lick trial’ (Fig. 1e). When calculating post-reward lick probability, if there was at least one lick within 32.4 cm after the reward location during a lap, this lap would be considered as a ‘lick trial’ (Extended Data Fig. 2c). The lick probability of a given bin, anticipatory lick probability, and post-reward lick probability were calculated as:

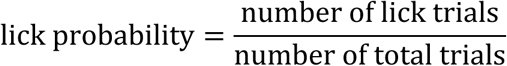

Post-reward lick latency was defined as the time interval between reward delivery and the first post-reward lick for each lap. Lick rate was defined as the number of licks divided by the dwell time within each spatial bin. To calculate post-reward lick rate, the lick rate of a given session was first averaged across laps. The post-reward lick rate was defined as the maximum lick rate within 21.6 cm after the reward location. All completed laps were included to plot and quantify the licking-related metrics.

#### Ca^2+^ signal extraction and processing

Images acquired across days were combined and registered using Suite2p (Python version, https://github.com/MouseLand/suite2p). Only data from the mice with stable FOVs across seven days were used for further processing. Representative FOVs in Fig. 1a and Extended Data Fig. 1a,b were average images of 500 frames acquired using the Z-project function in Fiji (version 2.16.0/1.54p). Image segmentation was first performed using Cellpose (version 3.0.10, in Python 3.9) to obtain regions of interest (ROIs). Frames were subsampled from all sessions using custom codes in Python and manually examined by the experimenter to ensure the quality of longitudinal tracking of individual neurons. A further quality control using principal component analysis (PCA) from randomly sampled continuous 60 frames per 500 frames throughout the longitudinal imaging was used to check for any functional contamination of anatomically identified ROIs by dendritic and somatic processes above or below the imaging plane. Only the ROIs that adequately and exclusively represented the same cell on each day were then selected for Ca^2+^ signal extraction using custom code in Python. All further analyses were conducted using custom code in MATLAB (version 2024a). Fluorescence smoothed by a moving-average filter (window size = 5) was converted to ΔF/F as:

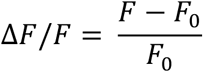

where F_0_ is the mode of the distribution of binned F within a 60-s moving window. The baseline noise threshold was calculated as follows. μ_0_ was the peak of the distribution of binned ΔF/F. A normal distribution was fitted to the binned data combining all values in X and 2μ_0_ – X, where X is all ΔF/F values below μ_0_, the peak of the distribution of binned ΔF/F. The baseline noise threshold was defined as μ + 3σ, where μ and σ were the mean and standard deviation of the normal distribution, respectively. Significant Ca^2+^ signals that exceeded this threshold for three continuous frames were considered as functional activity for further analysis. Locomotion-associated Ca^2+^ events were identified as the Ca^2+^ events that contained at least three frames recorded during the animal’s running (velocity > 2cm/s). All completed laps during imaging were used for further image analysis.

#### Quantification of CA1 activity during locomotion

Only locomotion-associated Ca^2+^ events were included in this analysis. The area under the curve (AUC) of a given event was calculated from its start to its end, where ΔF/F went above and below the baseline noise threshold respectively. To quantify activity level of each session, AUC and event numbers were normalized to the animal’s total running time in that session.

#### PC identification

PCs were detected as previously described^34,38,41^. Spatial maps of denoised neuronal activity (locomotion-associated transients above noise threshold described above) for each cell were constructed by evenly dividing the 180-cm track into 50 spatial bins (3.6 cm per bin). For each spatial bin, for every lap, the average ΔF/F was calculated using frames associated with locomotion (velocity >2 cm/s).

Each cell was assessed if it provided significant spatial information (SI) about the linear track location. SI for each cell was calculated as:

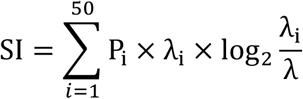

This was compared with SI of 500 shuffled trials of spatially binned activity for the same ROI, where each shuffle was performed by circularly shifting the ΔF/F values by at least 501 frames and then dividing the ΔF/F values into six roughly equal-sized chunks and permuting their order^34,66^. If the SI of the ROI exceeded 95% confidence interval determined by the shuffled trials, it was considered to have significant SI.

If a ROI had significant SI, we identified firing laps as laps with significant Ca^2+^ events whose peak was within a ±45-cm window of the average peak location for the session. The onset lap was determined as the first instance where at least three of five laps were firing laps. The reliability of putative PF activity was calculated as:

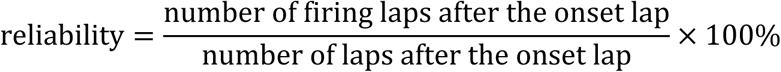

A cell was determined to have PF activity if (1) It had significant SI, (2) it had an onset lap (3/5 laps firing); and (3) the reliability from onset to end of session was at least 30%.

The width of PF activity in each firing lap was defined as the distance that the strongest Ca^2+^ event in the PF location spanned. The amplitude of PF activity in each firing lap was defined as the ΔF/F value of that the strongest Ca^2+^ event in the PF location. The coding properties of PCs for each session were averaged across firing laps (Extended Data Fig. 4a–e).

#### BTSP event identification

We searched for BTSP events in all cells as previously described^34,41^. A putative BTSP event was identified using the following criteria: (1) it was a strong Ca^2+^ event with an amplitude in the top 20th percentile of the event amplitudes of all firing laps in that cell for a given session; also, its amplitude was ≥ 1 ΔF/F; (2) only the laps with a peak ΔF/F within a ±45-cm window of the putative BTSP event peak location were considered as firing laps for further BTSP analysis; at least three out of five subsequent laps were firing laps; (4) the center of mass (COM) of the average activity in these firing laps showed a negative shift compared to the putative BTSP event location. The COM of neuronal activity was calculated as previously described^66^. The spatial activity map was transformed to polar coordinates, where θ denoted the bin location and r denoted the average ΔF/F in that bin. The center of mass of these points was calculated in a two-dimensional plane. Its angle was transformed back to the linear track, defined as the COM location. There could be multiple Ca^2+^ events meeting these criteria in a session. The minimum amplitude of these events was defined as the threshold for putative plateau potentials. The first Ca^2+^ event in a session that fulfilled these criteria was identified as a putative BTSP event. All the Ca^2+^ events whose amplitude was not lower than the plateau threshold were identified as putative plateaus.

The amplitude of spatially modulated activity occurring within a ±45-cm window of the BTSP event location in pre- and post-BTSP laps was measured in Fig. 2c. The peak location of the average activity following the BTSP lap was compared to the BTSP event location in Fig. 2d. The animal’s velocity during plateau induction was calculated as the average velocity in a ±30-frame window of the putative plateau in Fig. 2e.

In Extended Data Fig. 6e, plateau rate was calculated as number of putative plateaus divided by total lap number. To measure the dispersion of plateaus in Extended Data Fig. 6f, the plateau locations (0–180 cm) were first transformed to radians (0–2π). For these locations in radians {θ_i_}, the mean resultant vector length was computed as

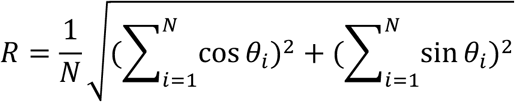

The circular variance (CV) of plateau locations was then defined as

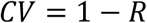

#### Determination of stable and unstable PCs

We tracked the PF activity of each cell from day 1 to day 7. On day n+1 (1 ≤ n ≤ 6), a PC would be considered stable if the cell was still identified as a PC and its PF location was within a ±30-cm window of the PF location on day n, as previously described^41^. In this case, subsequent comparisons of PFs would be made relative to the first-appearing PF location. If the PF shifts between day n and day n+1 exceeded 30 cm, or it became silent after having a PF, the cell would be considered unstable, and subsequent comparisons of PFs would be made relative to the newly appearing PF location (if any). On day n (1 ≤ n ≤ 7), the past PCs referred to the PCs that maintained their PF location established on previous days, and the new PCs referred to the PCs that had a new PF location on day n. On day n (1 ≤ n ≤ 7), the PC stability measured how long a PC had been stable at the PF location of day n. If a cell formed a new PF on day n, the PC stability was defined as one day. These analysis methods were applied in Fig. 3c–e and Extended Data Fig. 8a–d.

In Extended Data Fig. 8a, to characterize the decay dynamics of day-1- and day-2-appearing stable PCs, the data were fit to a double exponential function as (see ref. 34):

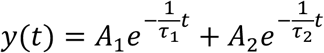

y(t) denotes the number of the PCs that remained stable since day 1 or 2. t is the time elapsed since day 1 or 2. A_1_ and *τ*1 represent the fast-decay component, and A_2_ and *τ*2 represent the slow-decay component (Extended Data Fig. 8b). Sustained PCs were identified as the PCs that maintained stable PFs for at least three days^34^.

In Fig. 4b, to calculate the probability of having a stable PF on the next day for a given level of PC stability, only PF locations on two consecutive days were compared. If the PF shift did not exceed ±30 cm, the cell would be considered stable.

#### Population vector (PV) correlation

To compare the stability of CA1 spatial representations across days, we conducted PV correlation analysis^34^. Spatial maps of neuronal activity were constructed as described in PC identification but using raw ΔF/F without denoising. PVs were activity vectors in which each element represents session-wise average activity of a neuron in a spatial bin. For each spatial bin, Pearson correlation coefficients were computed between PVs across days, producing a 7 × 7 matrix. For each animal, the data were averaged across the 50 spatial bins to generate a matrix M (also 7 × 7). To quantify cross-day correlation between genotypes, the correlation coefficient of k-day time interval (0 ≤ k ≤ 4) was calculated as (Fig. 3g):

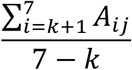

where *j* = *i* - *k*. A_ij_ is the element of matrix M in row *i* and column *j*.

#### Neural decoding

We used optimal linear estimation (OLE) method^67–69^ to decode the animal’s position on a circular track from neural population activity. The procedure for training a decoder and using it to decode neural activity is described below. Since the animal’s position is circular (ranging from 0 to 180 cm), we first convert it into an angular variable in radians, scaled from −π to π. Neural responses (raw ΔF/F without denoising) are binned across position and laps (3.6 cm per bin), resulting in a matrix R ∈ ℝ^M×N^, where M is the number of samples (spatial bins × laps) and N is the number of neurons. Each position θ ∈ [−π, π] is transformed into its circular components which serve as the target variables for linear regression:

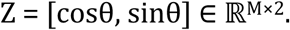

To normalize the neural responses, we subtract the mean activity of each neuron across samples:

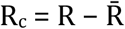

where R̄ ∈ ℝ^1×N^ is the mean response vector of each neuron.

We then compute the covariance matrix of the centered responses which captures the variability and shared noise across neurons:

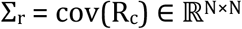

The cosine and sine target vectors are also mean-centered:

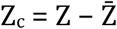

We then compute the cross-covariances between each position component and the neural responses:

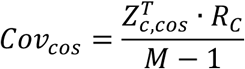

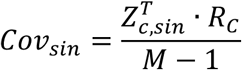

where Cov_cos_, Cov_sin_ ∈ ℝ^(1×N)^

To minimize the mean squared error, the optimal linear weights for decoding cosθ and sinθ are computed by:

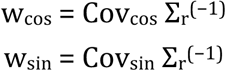

These weights linearly map neural activity to the corresponding circular component. The neural activity to be decoded, Rtest ∈ ℝ^Mtest×N^, is centered using the training mean:

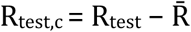

Decoded circular components are obtained by projecting the test responses onto the weight vectors:

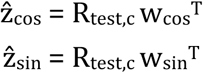

The decoded angle is reconstructed using the arctangent of the sine and cosine estimates:

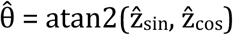

This gives the predicted angular position in radians, which can be transformed back to the decoded position on the linear track. The ground truth positions θ_true_ are also converted into an angular variable in radians, scaled from −π to π. The circular error between the predicted and actual positions is computed as:

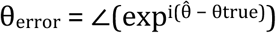

This ensures the circular error lies between −π and +π. The mean absolute error (converted to centimeters) was used to quantify decoding accuracy.

To validate decoding accuracy, we trained the decoder with data of the odd laps and used it to decode neural activity in the even laps of the same session (Extended Data Fig. 9a,b). Spatially binned neural activity (raw ΔF/F without denoising) was used for training. The neural response of each frame which was associated with locomotion in the even laps was decoded and generated a predicted position and decoding error (Extended Data Fig. 9a,b).

To compare the similarity of CA1 spatial representations across days, the decoder was trained using data from day i (1 ≤ i ≤ 7) and tested using neural activity from day j (1 ≤ j ≤ 7) (Fig. 3h,i and Extended Data Fig. 9c,d). For each pair of decoding, 150 randomly selected cells were used for training and testing, and this process was repeated 100 times. For each iteration, spatially binned neural activity (raw ΔF/F without denoising) was used for training. A time bin of 10 frames (∼0.333 s) was applied for testing. Decoding performance of each time bin was measured by the mean of errors (Extended Data Fig. 9c,d). For each animal, the decoding error across days was an average across time bins and iterations. When quantifying cross-day decoding errors between genotypes, we subtracted the same-day decoding error from the decoding error matrix, producing matrix N. The decoding error change of k-day time interval (0 ≤ k ≤ 4) was calculated as (Fig. 3i):

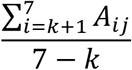

where j = i-k. A_ij_ is the element of matrix N in row i and column j.

#### Determination of in-field and out-of-field BTSP events

This analysis was only applied to the cells that had PFs on two consecutive days, day n (1 ≤ n ≤ 6) and day n+1, and had BTSP events at least on day n+1. The location of the BTSP event on day n+1 was compared with the PF location on day n. If the BTSP event on day n+1 fell into a ±30-cm window of the PF location on day n, it would be considered as an in-field BTSP event (Fig. 4f). Otherwise, it would be considered as an out-of-field BTSP event (Fig. 4g).

#### Characterization of residual activity

The analyses of residual activity were only applied to the cells that had PFs on two consecutive days, day n (1 ≤ n ≤ 6) and day n+1, and had BTSP events at least on day n+1. Only the five laps prior to the BTSP lap were used for these analyses. If there were fewer than five laps before the BTSP lap, all available laps were included. To present the process of PF re-emergence, the last three firing laps (see PC identification) on day n were included, and the laps on day n+1 were aligned to the BTSP lap (Fig. 4c). On day n+1, the amplitude of PF activity denoted the amplitude of firing laps. If a lap surrounding the BTSP lap was not identified as a firing lap, its amplitude was defined as 0. All available BTSP events were combined in Fig. 4c.

The in-field residual activity on day n+1 was defined as locomotion-associated Ca^2+^ events that occurred before the BTSP lap within a ±28.8-cm window of the PF location on day n. The amplitude of the in-field residual activity was averaged across the laps included in this analysis (Fig. 4e,f). If no significant in-field Ca^2+^ events occurred in a lap, the amplitude of that lap was defined as 0. Similarly, the amplitude of the out-of-field pre-BTSP activity was calculated for locomotion-associated Ca^2+^ events occurring outside the ±28.8-cm window of the PF location on day n. The selectivity index of residual activity was calculated as (Fig. 4h):

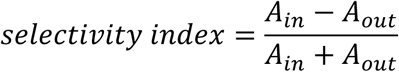

A_in_ is the average amplitude of in-field residual activity, and A_out_ is the average amplitude of out-of-field pre-BTSP activity.

To measure the similarity between residual activity and previous PF, we calculated Pearson correlation coefficient between lap-wise activity on day n+1 (1×50 vector for each lap) and PF activity on day n that was also a 1×50 vector after averaging all firing laps (see PC identification) (Fig. 4i). All available BTSP events were combined in Fig. 4i. For animal-wise comparison, the activity of at most five laps before the BTSP lap was averaged across laps to produce a 1×50 vector, which was used for Pearson correlation calculation as described above (Fig. 4j).

In Fig. 4b,d,e,g,h,j, PC stability was calculated as described in Determination of stable and unstable PCs.

### PC dynamics model

We modeled the across-day dynamics of hippocampal CA1 PFs using an ensembled synaptic weight model, allowing each neuron to maintain multiple distinct PFs (up to *n* = 5 per neuron). Each potential PF *k* of neuron *i* on session (day) *t* is characterized by an ensembled synaptic weight 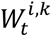, representing the average synaptic weights of that field. These weights evolve across days following an activity-dependent exponential decay rule. After each daily session, the weight of each field is updated as follows:

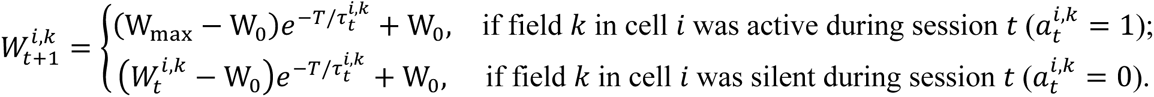

Here W_0_ is the baseline synaptic weight (minimum strength), W_max:_ is the maximum weight, and 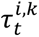 is the decay time constant governing how quickly the weight decays over the inter-session interval *T* (the time between sessions). If a PF was active on day 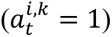, we assumed that its weight was potentiated to the maximal value W_89:_ during the session via BTSP and then decays from that peak value. In contrast, a silent field’s weight decays from its current value back toward W_max_. This captures the intuition that regularly reactivated PFs retain high synaptic weights through repetitive BTSP events, while silent fields gradually lose strength across days.

PC activity on each day is modeled as a two-step stochastic process. First, each neuron i is designated as either active or silent in session *t* using a binary variable 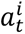. The probability of neuron i being active on day *t* is given by:

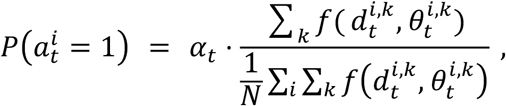

where *N* is the total number of neurons and *α_t_* is a scaling factor that controls the overall fraction of active cells. The depolarization activity level 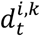 is computed as:

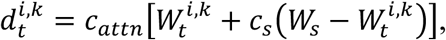

combining synaptic input proportional to the weight 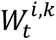. with a spontaneous activity component 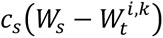, scaled by the attenuation factor *c_attn_*. We also include global feedback inhibition that raises the threshold for plateau potential firing as overall network activity increases. Specifically, the global inhibitory feedback term scales with averaged depolarization above baseline across all fields:

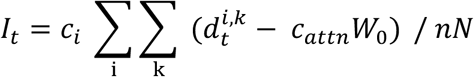

and the threshold of firing plateau for each field is set as 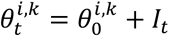. Baseline thresholds 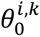 are heterogenous across all fields, drawn independently from a piecewise uniform distribution:

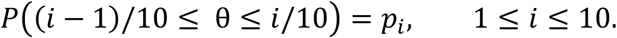

The function

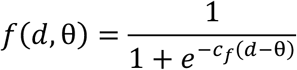

is a logistic sigmoid activation function (with steepness *c_f_*) that converts the depolarization *d* into an activation propensity given threshold θ. The expected fraction of active cells *α_t_* scales with overall network activity as

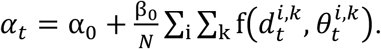

Thus, when the population is more active (e.g. under the RTT condition with higher spontaneous drive *c*_2_), *α*_+_ increases and a larger fraction of neurons will be PCs. Finally, we determine each neuron’s state for each day *t* by sampling all 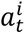 from Bernoulli distribution parametrized by the event probability 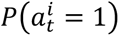.

Second, we assume that if cell *i* is active on day *t* 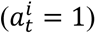, it will express exactly one of its *n* potential PFs. We denote the activity of a specific field by a one-hot categorical variable 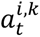, which equals 1 for the selected field *k* and 0 for all other fields of that neuron. The probability of selecting field *k* (conditional on cell *i* being active) is defined by a softmax function over the field’s activation propensities:

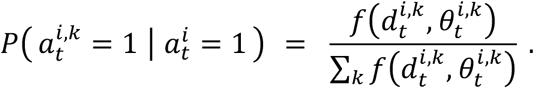

This stochastic field-selection mechanism ensures that only one PF per active cell is expressed in any session, while allowing different fields of the same neuron to appear on different days. The factor *f*(*d*, θ) biases selection toward fields with higher depolarization 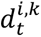 but retains variability so that even weaker fields can occasionally be expressed. Note that a field is active 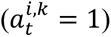 if and only if its neuron is active and it was the field selected for that neuron.

To incorporate long-term consolidation effects, we also introduce a progressively stabilized synapse model inspired by the cascade model of synaptic consolidation^70^. Specifically, each time a PF is formed or reconstituted, its decay time constant is increased, making that field more stable across days. This stabilization follows the rule:

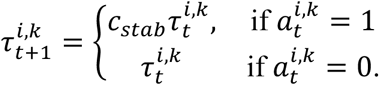

with initially 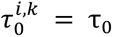 for all fields. Repeated reactivation progressively lengthens the decay time constant, mimicking a cascade of synaptic consolidation, whereas silent fields retain the same time constant and remain more transient.

Model parameters are fitted to reproduce key statistics of experimentally observed PC behavior. The fitting procedure minimize a loss function comparing the model output to experiments on the following three metrics: (i) the distribution of active days per cell (Extended Data Fig. 10a), (ii) the day-by-day evolution of active PC counts (Extended Data Fig. 10b), and (iii) the history-dependent probability of PF formation (Extended Data Fig. 10c) and PF reappearance after one inactive day (Extended Data Fig. 10d). We also introduce a regularization term constraining the depolarization level 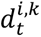 remained close between WT and RTT conditions. Model parameters were optimized using a genetic algorithm (MATLAB Global Optimization Toolbox), minimizing the maximum loss across both WT and RTT conditions (plus the regularization term), so that both conditions are fitted simultaneously.

Notably, the only parameter that differs between WT and RTT conditions is the parameter *c*_2_ scaling the spontaneous activity (set to 0 for WT versus 0.398 for RTT); all other parameters are identical across two conditions. The final fitted parameters are: *n* = 5, *T* = 23 ℎ*ours*, *τ*_0_ = 11.85 ℎ*ours*, *W*_0_ = 0.14, W_max_ = 1, *c*_attn_ = 0.37, *W*_s_ = 0.79, *c*_i_ = 0.76, α_0_ = 0.12, β_0_ = 0.90, *c_f_* = 26.47, *c_stab_* = 1.114, and *p*_1_, *p*_2_, ⋯, *p*_10_ = 0.007, 0.191, 0.194, 0.120, 0.148, 0.094, 0.076, 0.032, 0.090, 0.048. This single set of parameters was used to simulate both WT and RTT conditions (with only *c*_s_ differing as noted above) and was sufficient to capture the experimentally observed PF dynamics in each condition. The simulation is repeated for 10 times for each condition.

To simulate the experimental observations, the analysis methods of modeled data resembled those describe above for experimental data (Fig. 5c–h and Extended Data Fig. 10a–e). We assume that all the PFs in the model are driven by plateau potentials and BTSP. For the model, only the PCs that maintained the same PF location were considered as stable ones and no jittering was tolerated for stable PCs. The in-field residual amplitude was defined as 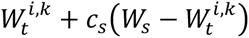 The out-of-field pre-BTSP activity amplitude was calculated in the same way but was the maximum activity level across the four out-of-field PF locations.

For the simulation that manipulated only 50% of the cells under RTT-like conditions, we use the same PC dynamics model as described above. The only parameter that differs between WT and RTT conditions is the parameter *c*_2_ scaling the spontaneous activity (set to 0 for WT versus 0.404 for RTT); all other parameters are identical across two conditions. The final fitted parameters are: *n* = 5, *T* = 23 ℎ*ours*, *τ*_0_ = 10.13 ℎ*ours*, *W*_0_ = 0.18, W_max_ = 1, *c*_attn_ = 0.41, *W_s_* = 1.00, *c_i_* = 0.83, α_0_ = 0.14, β_0_ = 0.67, *c_f_* = 28.77, *c_stap_* = 1.164, and *p*_1_, *p*_2_, ⋯, *p*_10_ = 0.001, 0.192, 0.182, 0.137, 0.096, 0.060, 0.115, 0.0114, 0.029, 0.074. The simulation is repeated for 10 times for each condition.

## Statistical methods

The exact sample size (*n*) of each experiment and analysis is reported in the figure legends or in the main text. No power analysis was used to predetermine sample sizes, but our sample sizes are comparable to those reported in previous publications that performed similar behavioral task and population imaging^25,32,38,41,47^. Data are shown as mean ± SEM unless otherwise specified. All the statistical tests were conducted using built-in and custom functions in MATLAB.

## Contributions

J.C.M. and H.Y.Z. conceived the project. T.R.Z. performed the experiments supervised by S.P.V.. G.L. performed the computational modeling. T.R.Z. and S.P.V. analyzed the experimental data and wrote the manuscript with inputs from all authors.

## Acknowledgements

We would like to thank Dr. Raymond Chitwood for technical support and other members of the Magee lab and Zoghbi lab for helpful discussions. We thank Yaling Sun for mouse colony management. We thank Dr. Rachel Arey, Dr. Barna Dudok, Dr. Daoyun Ji, and Yan Li for providing useful comments on the manuscript. This work was funded by Howard Hughes Medical Institute (J.C.M. and H.Y.Z.), the National Institute of Neurological Disorders and Stroke (NINDS) (5R01NS057819 to H.Y.Z.) and the Cullen Foundation (J.C.M.).

## Data availability

The data supporting this study’s findings are available from the corresponding authors upon request.

## Code availability

Scripts that support the results of this study will be available on GitHub before publication and can be requested from the corresponding authors till then.

---

**Extended Data Fig. 1.**
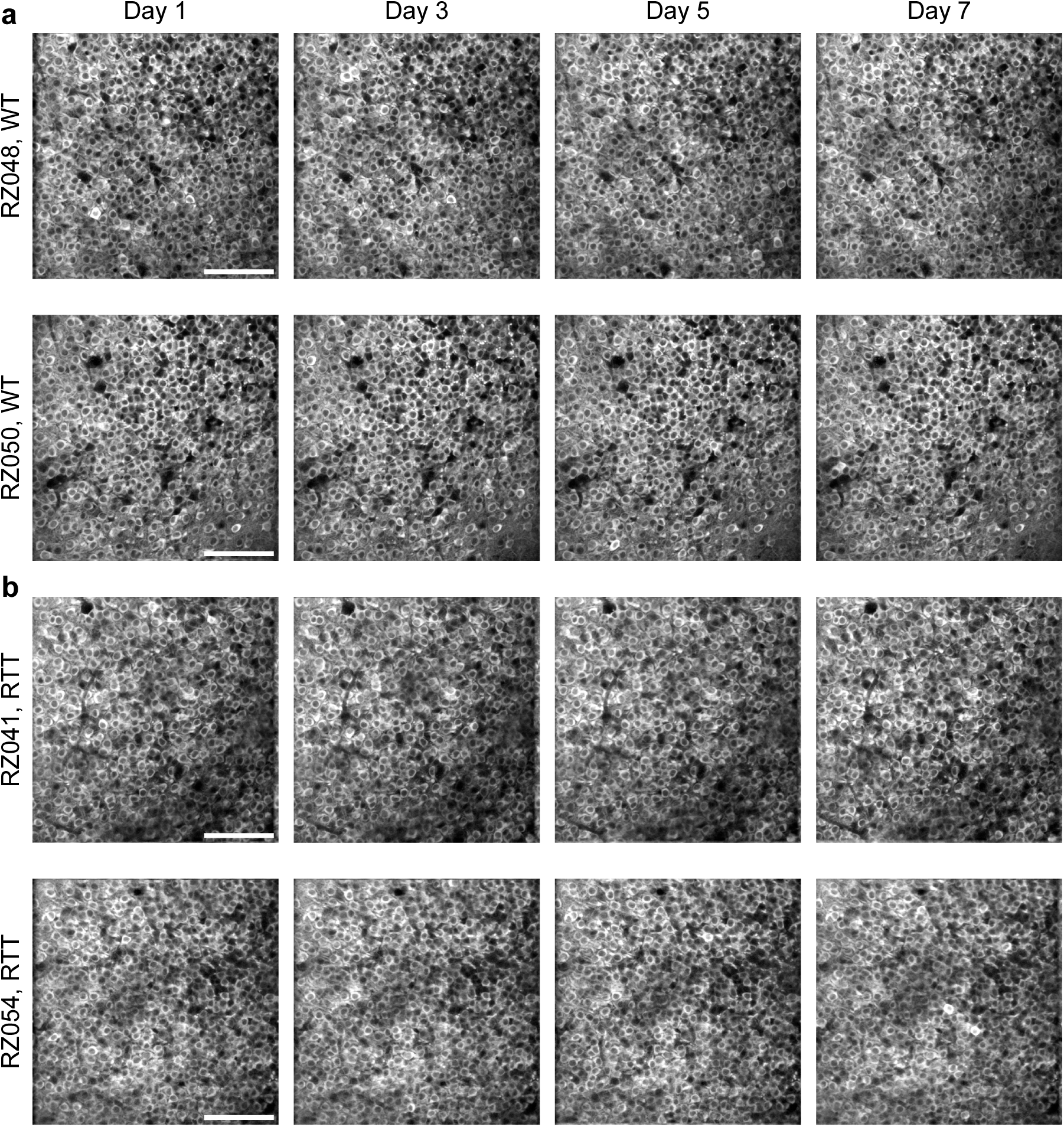
Longitudinal imaging of the same population of CA1 pyramidal neurons over multiple days. **a,b,** Representative field of views (FOVs) of two WT mice (**a**) and two RTT mice (**b**) showing stable access to the same neuronal population across seven days. Each FOV is an average z-projection of 500 consecutive frames that were randomly selected. Scale bar, 100 µm.

**Extended Data Fig. 2.**
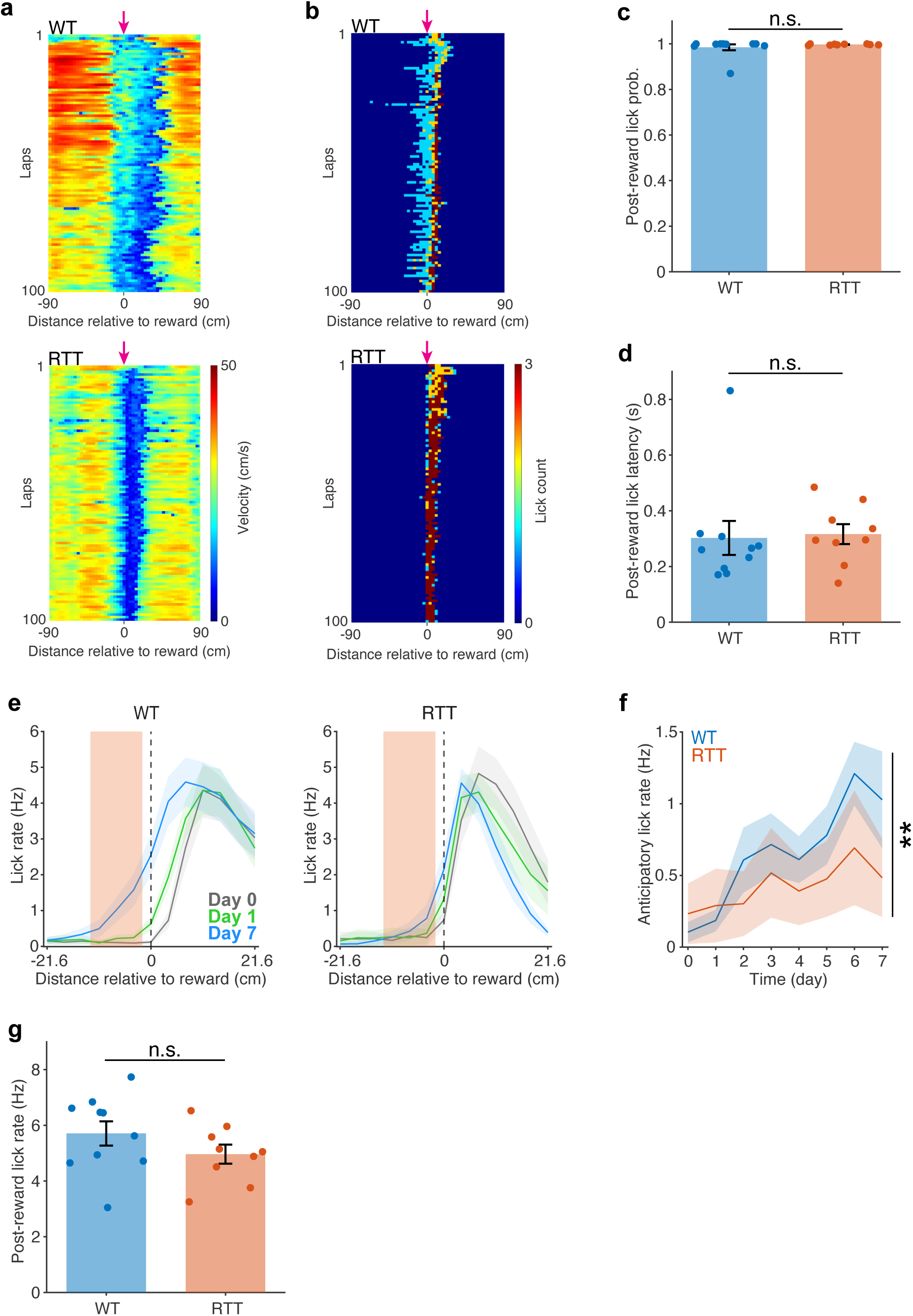
Additional measures of adaptive behaviors. **a,** Running velocity across laps of a WT mouse (top) and a RTT mouse (bottom). The magenta arrow indicates the reward location (**a**,**b**). **b,** Number of licks across laps of a WT mouse (top) and a RTT mouse (bottom). **c,** Lick probability after reward delivery in WT and RTT mice (see Methods). Data of individual animals are averaged across days (Day 1–7). Two-sample *t*-test, *p* = 0.386. n.s., *p* > 0.05. **d**, Lick latency to reward delivery in WT and RTT mice (see Methods). Data of individual animals are averaged across days (Day 1–7). Two-sample *t*-test, *p* = 0.854. n.s., *p* > 0.05. **e,** Lick rate around the reward location in WT (left) and RTT (right) mice during the entire sessions on day 0, day 1, and day 7. Orange boxes indicate the anticipatory zone, in which the lick rate is measured for comparison in **f**. **f,** Lick rate in the anticipatory zone versus time. Linear mixed-effects model, time: *p* = 0.022; genotype × time interaction: *p* = 0.002. **, genotype × time interaction *p* < 0.01. **g**, Peak lick rate after reward delivery in WT and RTT mice (see Methods). Data of individual animals are averaged across days (Day 1–7). Two-sample *t*-test, *p* = 0.2037. n.s., *p* > 0.05. **c**–**g,** WT, *n* = 10 mice; RTT, *n* = 9 mice. Data across animals are presented as mean ± SEM.

**Extended Data Fig. 3.**
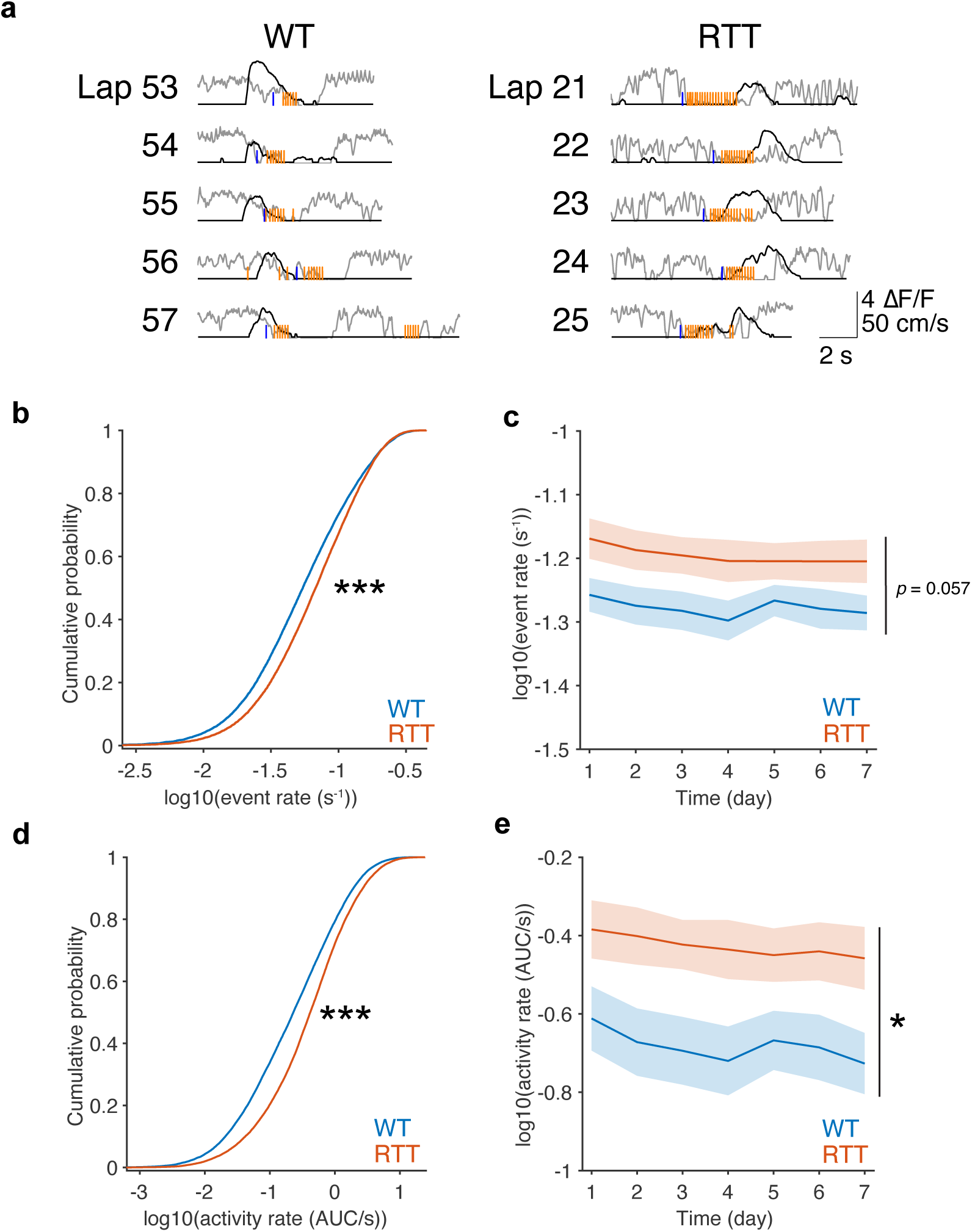
RTT mice exhibit increased activity in CA1 area of the hippocampus. **a,** Example traces of neuronal activity (ΔF/F, black) of two PCs and corresponding animal behavior, including velocity (gray) and licks (orange). Each row represents a lap. The blue lines indicate reward delivery. Left: WT; right: RTT. **b,** Cumulative probability of logarithmic Ca^2+^ event rate (log10(number of events/s), see Methods) in WT and RTT mice. Data from all cells and across seven days are combined. WT, *n* = 24486 cell × day; RTT, *n* = 22358 cell × day. Kolmogorov–Smirnov test, *p* = 1.670 × 10^−97^. ***, *p* < 0.001. **c,** Animal-wise comparison of logarithmic Ca^2+^ event rate across days. Two-way repeated-measures ANOVA, *p* = 0.057; time: *p* = 2.218 × 10^−21^. **d,** Same as **b**, but for logarithmic activity rate (log10(AUC/s), see Methods). Kolmogorov–Smirnov test, *p* = 6.162 × 10^−224^. ***, *p* < 0.001. **e,** Same as **c**, but for logarithmic Ca^2+^ activity rate. Two-way repeated-measures ANOVA with the Greenhouse-Geisser correction, genotype: *p* = 0.030; time: *p* = 7.100 × 10^−9^. *, genotype *p* < 0.05. **c,e,** WT, *n* = 10 mice; RTT, *n* = 9 mice. Data are presented as mean ± SEM.

**Extended Data Fig. 4.**
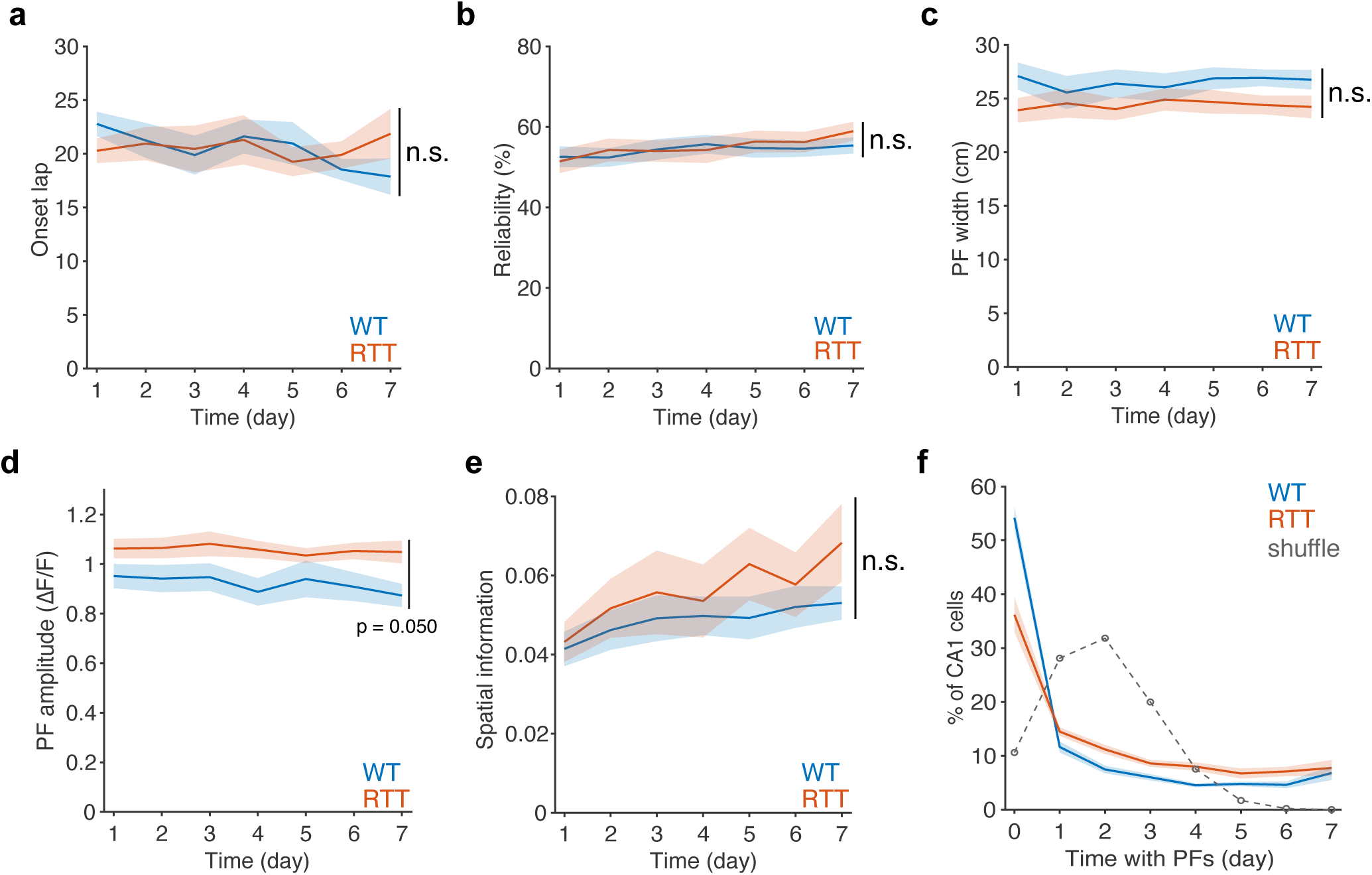
Comparison of PC properties between RTT and WT mice. **a,** Average onset lap number of PCs across days. Two-way repeated-measures ANOVA, genotype: *p* = 0.931; time: *p* = 1.043 × 10^−13^. n.s., genotype *p* > 0.05. **b,** Average reliability of PF firing across days. Two-way repeated-measures ANOVA, genotype: *p* = 0.806; time: *p* = 2.631 × 10^−17^. n.s., genotype *p* > 0.05. **c,** Average PF width (in centimeter) across days. Two-way repeated-measures ANOVA, genotype: *p* = 0.154; time: *p* = 6.072 × 10^−18^. n.s., genotype *p* > 0.05. **d,** Average PF amplitude across days. Two-way repeated-measures ANOVA, genotype: *p* = 0.050; time: *p* = 3.903 × 10^−16^. **e,** Average spatial information of PCs across days. Two-way repeated-measures ANOVA, genotype: *p* = 0.409; time: *p* = 1.238 × 10^−9^. n.s., genotype *p* > 0.05. **f,** Distribution of total number of days with PF activity. The solid lines show long-tailed distributions in both genotypes. The gray dashed line indicates data from a random process. **a**–**f,** WT, *n* = 10 mice; RTT, *n* = 9 mice. Data are presented as mean ± SEM. The analyses of PC coding properties are described in Methods.

**Extended Data Fig. 5.**
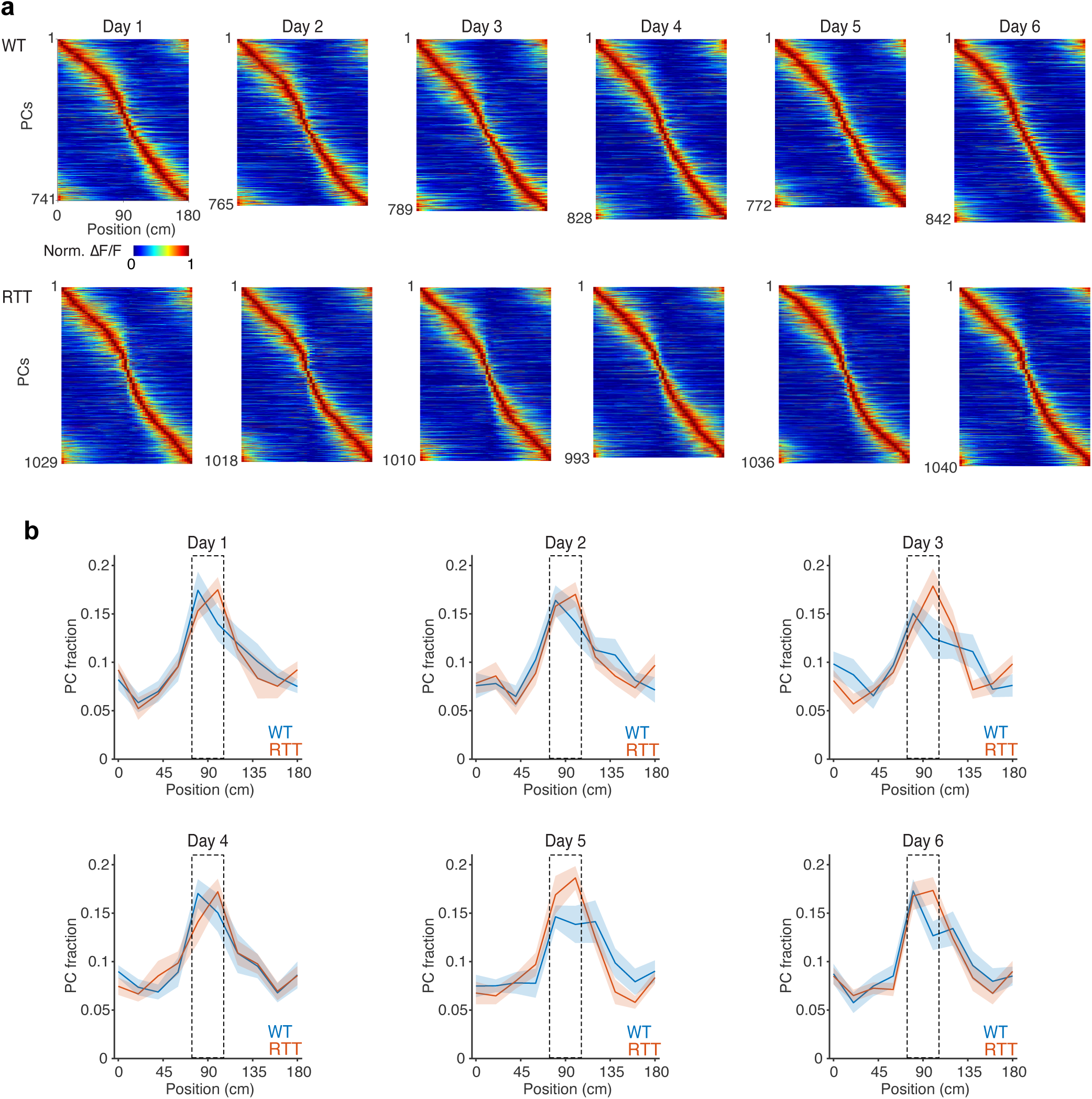
Preserved overrepresentations of the reward zone in RTT mice. **a,** Peak-normalized activity (ΔF/F) of all PCs on day 1–6 in WT mice (top) and RTT mice (bottom). PCs are combined across animals and sorted by their PF locations. **b**, Normalized distribution of PF locations on day 1–6. The box indicates the reward zone. Bin = 18 cm. WT, *n* = 10 mice; RTT, *n* = 9 mice. Data are presented as mean ± SEM. Day 1, Chi-squared test, WT: df = 9, *p* = 4.413 × 10^−15^; RTT: df = 9, *p* = 1.144 × 10^−23^. Day 2, Chi-squared test, WT: df = 9, *p* = 4.411 × 10^−12^; RTT: df = 9, *p* = 4.129 × 10^−20^. Day 3, Chi-squared test, WT: df = 9, *p* = 7.418 × 10^−8^; RTT: df = 9, *p* = 1.144 × 10^−25^. Day 4, Chi-squared test, WT: df = 9, *p* = 3.254 × 10^− 14^; RTT: df = 9, *p* = 1.036 × 10^−15^. Day 5, Chi-squared test, WT: df = 9, *p* = 8.779 × 10^−10^; RTT: df = 9, *p* = 2.663 × 10^−35^. Day 6, Chi-squared test, WT: df = 9, *p* = 3.646 × 10^−14^; RTT: df = 9, *p* = 3.057 × 10^−26^.

**Extended Data Fig. 6.**
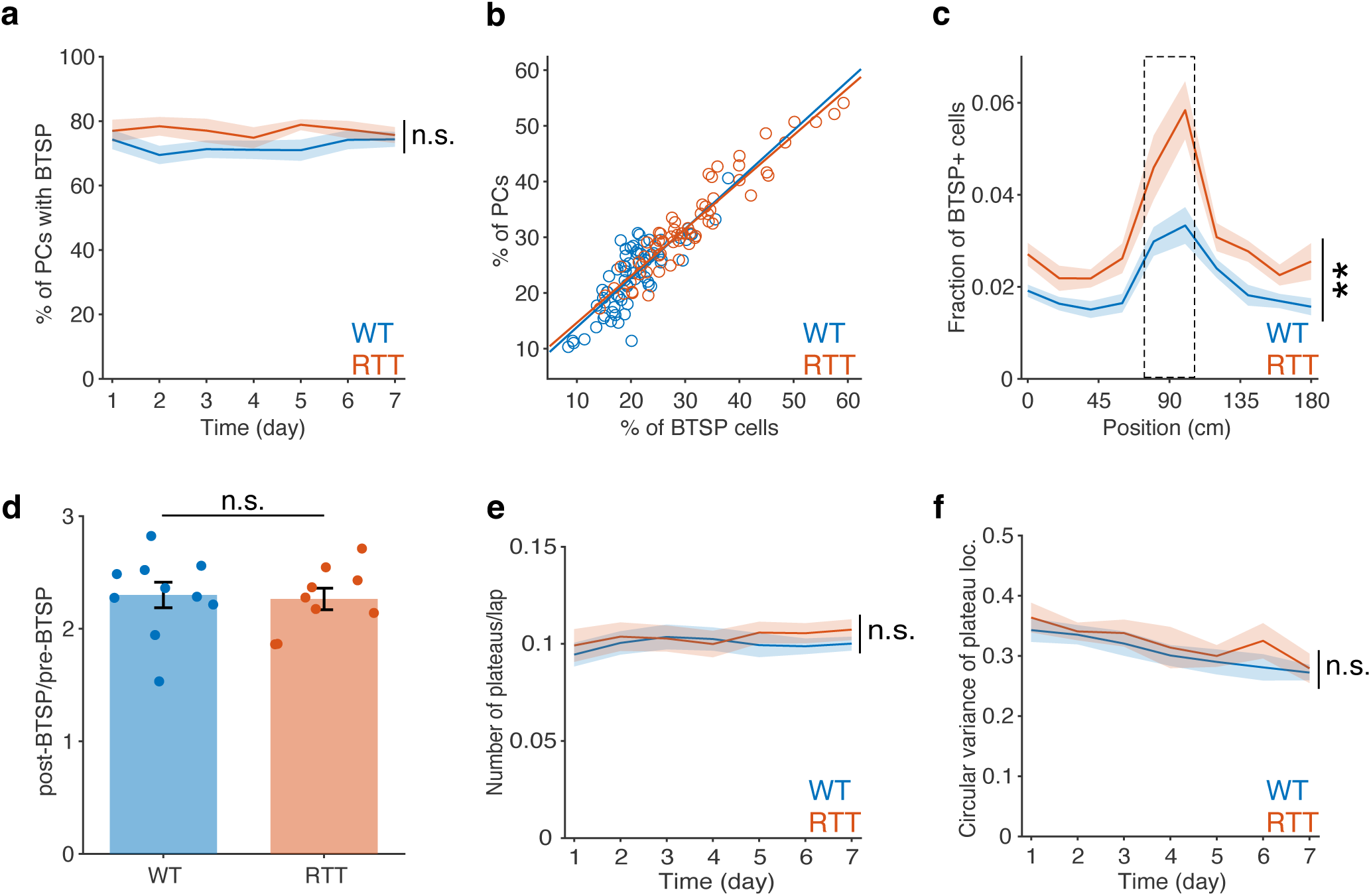
Properties of BTSP and plateau potentials remain intact in RTT mice. **a,** Percentage of PCs showing atleast one BTSP event among CA1 PCs across days. Two-way repeated-measures ANOVA, genotype: *p* = 0.142; time: *p* = 8.888 × 10^−20^. n.s., genotype *p* > 0.05. **b,** Percentage of PCs as a function of percentage of cells with BTSP events. Each dot represents an individual recording session. A linear regression model was fitted to the data. WT: R^2^ = 0.66116, *p* = 1.232 × 10^−17^, y = 0.88x + 5.01 (blue line). RTT: R^2^ = 0.90234, *p* = 1.641 × 10^−32^, y = 0.84x + 6.23 (orange line). **c,** Distribution of BTSP+ cells along the track (normalized to the total cell number). The box indicates the reward zone. Bin = 18 cm. Two-way repeated-measures ANOVA with the Greenhouse-Geisser correction, genotype: *p* = 0.007; bin: *p* = 3.416 × 10^−11^. **, genotype *p* < 0.01. **d,** Ratio of post-BTSP ΔF/F to pre-BTSP ΔF/F showing the abrupt increase in spatially modulated activity following BTSP induction, corresponding to Fig. 2c. Two-sample *t*-test, *p* = 0.812. n.s., *p* > 0.05. **e,** Plateau rate (number of plateaus per lap in BTSP identified cells, see Methods) across days. Two-way repeated-measures ANOVA with the Greenhouse-Geisser correction, genotype: *p* = 0.680; time: *p* = 6.913 × 10^−16^. n.s., genotype *p* > 0.05. **f,** Circular variance of plateau locations in BTSP identified cells on the belt (see Methods) across days. Two-way repeated-measures ANOVA, genotype: *p* = 0.508; time: *p* = 3.127 × 10^−14^. n.s., genotype *p* > 0.05. **a–f**, WT, *n* = 10 mice; RTT, *n* = 9 mice. a,c–f, Data are presented as mean ± SEM.

**Extended Data Fig. 7.**
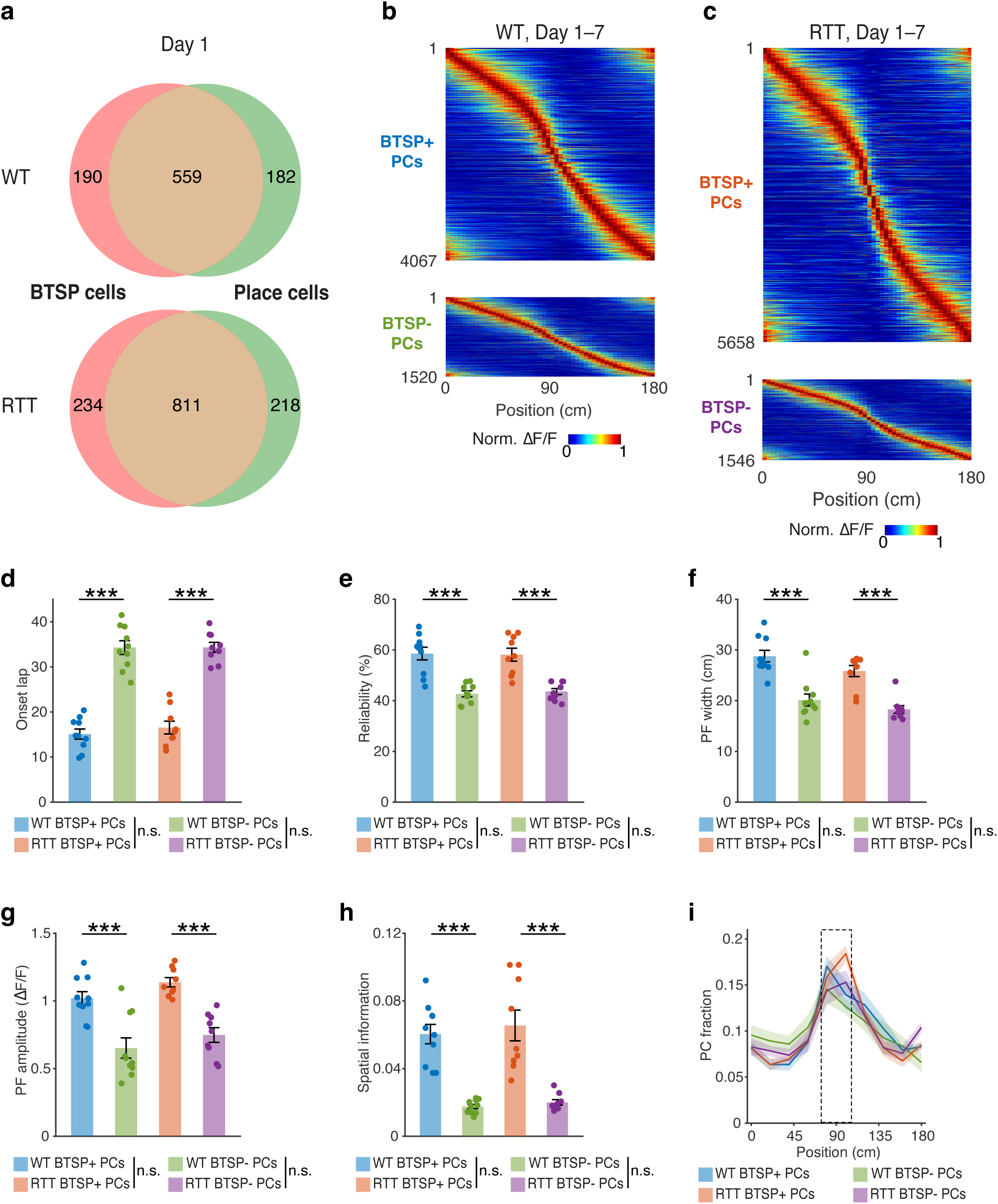
Comparison of coding properties between BTSP+ PCs and BTSP-PCs. **a,** Venn diagrams showing the overlap between BTSP+ cells and PCs on day 1 in WT mice (top) and RTT mice (bottom). Cells are combined across animals. **b**, Peak-normalized activity (ΔF/F) of BTSP+ PCs (top) and BTSP- PCs (bottom) in WT mice. PCs are combined across animals and sessions and sorted by their PF locations. **c**, Same as **b**, but for RTT mice. **d,** Average onset lap number of BTSP+ PCs and BTSP- PCs. One-way ANOVA, *p* = 2.345 × 10^−^14. **e,** Average PF firing reliability of BTSP+ PCs and BTSP- PCs. One-way ANOVA, *p* = 1.242 × 10^−7^. **f,** Average PF width (in centimeter) of BTSP+ PCs and BTSP- PCs. One-way ANOVA, *p* = 6.704 × 10^−8^. **g,** Average PF amplitude of BTSP+ PCs and BTSP- PCs. One-way ANOVA, *p* = 8.881 × 10^−7^. **h,** Average spatial information of BTSP+ PCs and BTSP- PCs. One-way ANOVA, *p* = 2.898 × 10^−8^. **i**, Normalized distribution of PF locations of BTSP+ PCs and BTSP- PCs. The box indicates the reward zone. Bin = 18 cm. Chi-squared test, WT BTSP+ PCs: df = 9, *p* = 6.106 × 10^−86^; WT BTSP-PCs: df = 9, *p* = 1.360 × 10^−10^; RTT BTSP+ PCs: df = 9, *p* = 1.961 × 10^−171^; RTT BTSP- PCs: df = 9, *p* = 2.541 × 10^−20^. **d–h,** Statistical significance between groups is assessed using Tukey’s multiple-comparisons test following one-way ANOVA. n.s., *p* > 0.05; ***, *p* < 0.001. **d–i,** WT, *n* = 10 mice; RTT, *n* = 9 mice. Data of individual animals are averaged across days. Data are presented as mean ± SEM.

**Extended Data Fig. 8.**
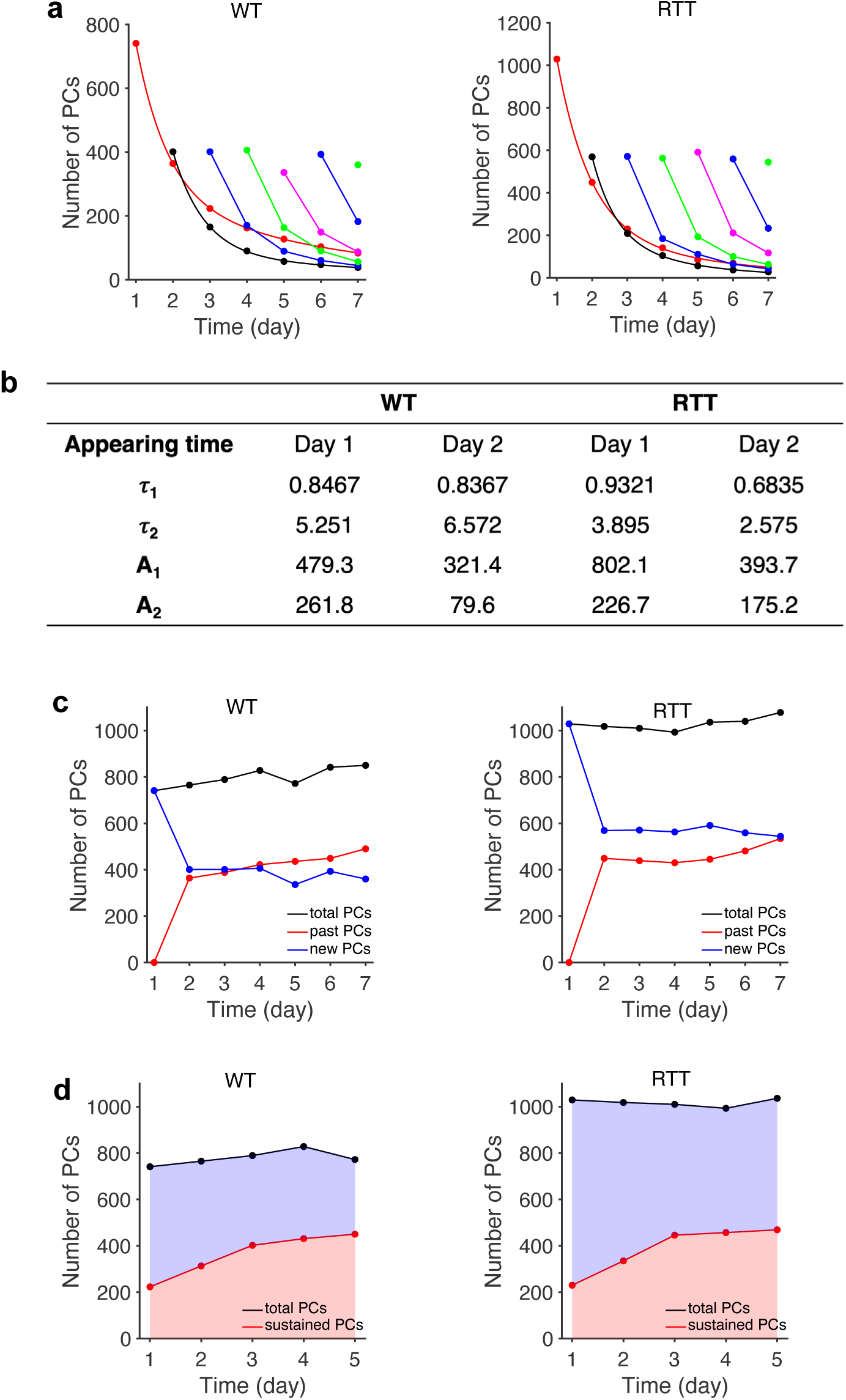
Development of stable memory representations is disrupted in RTT mice. **a,** Dynamics of the number of PCs that first appear on day 1–7 and maintain a consistent PF location on subsequent days in WT (left) and RTT (right) mice (see Methods). Colors represent when PCs first appear. Red, day 1; black, day 2; blue (left), day 3; light green, day 4; magenta, day 5; blue (right), day 6; dark green, day 7. Double exponential functions indicated by the smooth lines are fitted to the data of PCs first appearing and day 1 (red) and day 2 (black) (see Methods). **b,** Parameters of the double exponential functions for PCs first appearing and day 1 and day 2 (see Methods). **c,** Number of past PCs (maintaining previously established PFs) and new PCs (form PFs at new locations) versus time in WT (left) and RTT (right) mice, related to Fig. 3c. **d,** Number of sustained PCs displaying enhanced stability (see Methods and ref. 34) versus time in WT (left) and RTT (right) mice. **a**–**d,** PCs are combined across animals.

**Extended Data Fig. 9.**
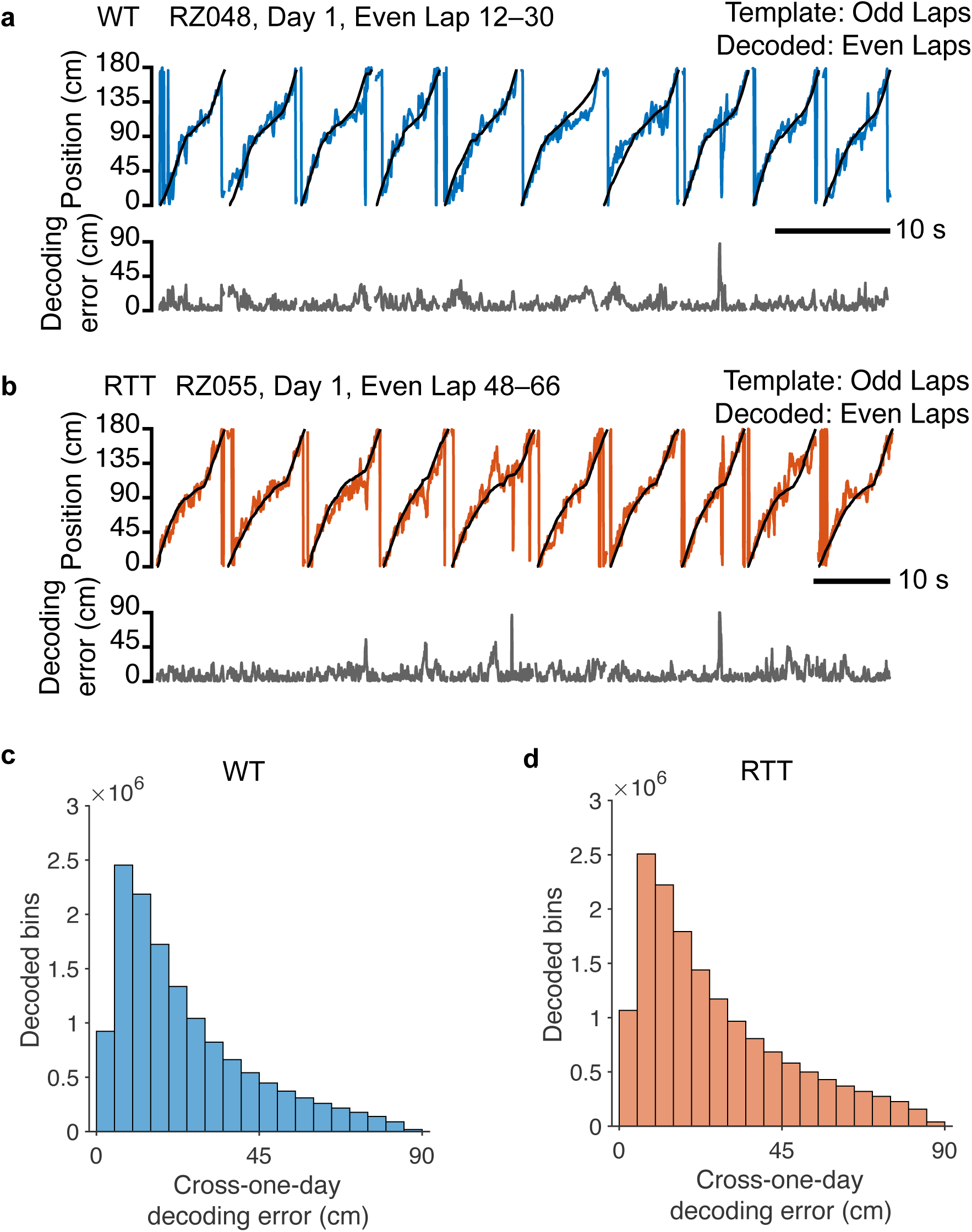
Same-day and cross-day linear decoding of population activity. **a,** Real-time decoding performance using the optimal linear estimator (OLE) method trained with the odd laps and tested with the even laps from one given session (see Methods) in a WT mouse. Top: blue, the animal’s actual position; black, decoded position. Bottom: decoding error. **b,** Same as **a**, but for a RTT mouse. Top: orange, the animal’s actual position; black, decoded position. Bottom: decoding error. **c,** A representative distribution of cross-day decoding errors (see Methods) for a WT mouse. The decoder is trained with day-1 data and tested with day-2 data from randomly selected 150 cells and iterated for 100 times. Decoded bin = 10 frames (∼0.333 s). *n* = 2.305 × 10^5^ bins. **d,** Same as **c**, but for a RTT mouse. *n* = 2.579 × 10^5^ bins.

**Extended Data Fig. 10.**
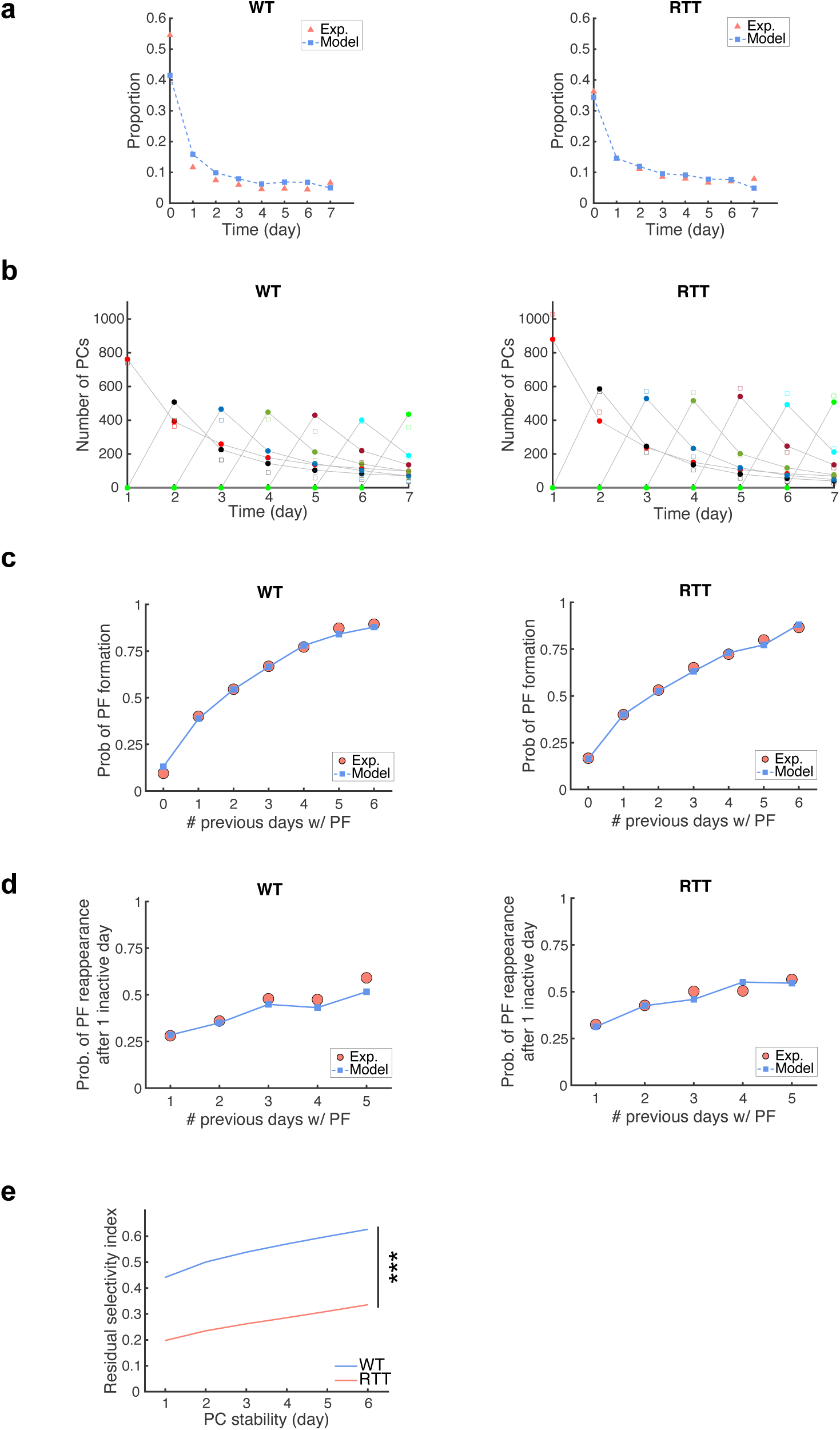
Fitting procedure of the PC dynamics model. **a–d,** The model parameters are fitted to the key statistics of experimentally observed PCs dynamics as follows (see Methods). **a**, the distribution of the number of active days with PFs. **b**, the day-by-day evolution of stable PC counts. **c**, the history-dependent PF formation probability. **d**, the history-dependent probability of PF reappearance after one inactive day. Both WT-like and RTT-like conditions are fitted at the same time to minimize the loss function. The experimental data are depicted by triangles (**a**), squares (**b**), or circles (**c,d**). The modeled data are depicted by squares (**a,c,d**) or circles (**b**). **e,** The PF dynamics model recapitulates the diminished selectivity of residual activity under the RTT-like conditions. Two-way repeated-measures ANOVA with the Greenhouse-Geisser correction, genotype: *p* = 4 × 10^−42^. ***, genotype *p* < 0.001. WT-like condition, *n* = 10 simulations; RTT-like condition, *n* = 10 simulations. Data are presented as mean ± SEM. The error bars are not visible because of the low variance.

**Extended Data Fig. 11.**
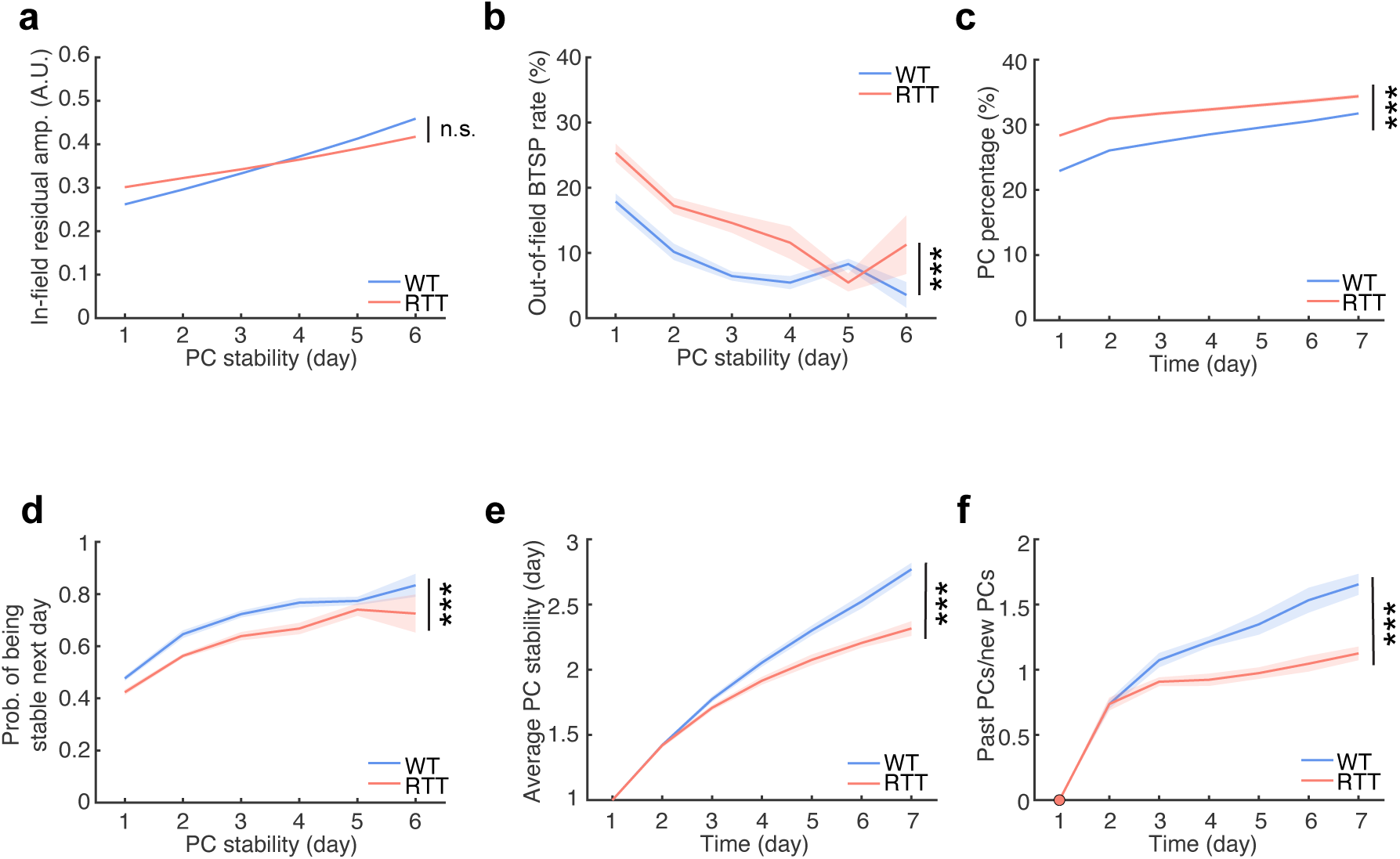
A computational model with altered residual selectivity in half the neurons recapitulates RTT PC dynamics. **a**, The in-field residual amplitude stays comparable between the WT- and RTT-like conditions in this model. Two-way repeated-measures ANOVA with the Greenhouse-Geisser correction, genotype: *p* = 0.09. n.s., genotype *p* > 0.05. **b,c,d,** The PF dynamics model recapitulates the out-of-field BTSP probability (**b**), percentage of PCs (**c**), and experience-dependent PC stabilization (**d**) under the WT- and RTT-like conditions. Two-way repeated-measures ANOVA with the Greenhouse-Geisser correction. **b**, genotype: *p* = 9 × 10^−5^. **c**, genotype: *p* = 1 × 10^−10^. **d**, genotype: *p* = 5 × 10^−4^. ***, genotype *p* < 0.001. **e,f,** The PF dynamics model recapitulates the development of stable CA1 representations with task experience and the divergent responses between the WT- and RTT-like conditions. **e**, Average PC stability measuring the number of days during which the PC has maintained the same PF location. Two-way repeated-measures ANOVA with the Greenhouse-Geisser correction, genotype: *p* = 5 × 10^−5^. ***, genotype *p* < 0.001. **f**, Ratio of past PCs (maintaining previously established PFs) to new PCs (form PFs at new locations). Two-way repeated-measures ANOVA with the Greenhouse-Geisser correction, genotype: *p* = 2 × 10^−6^. ***, genotype *p* < 0.001. **a**–**f,** WT-like condition, *n* = 10 simulations; RTT-like condition, *n* = 10 simulations. Data are presented as mean ± SEM. The error bars in **a** and **c** are not visible because of the low variance.

